# Riboflavin Directly Mediates the Dealkylation by Microbial Cytochrome P450 Monooxygeneses

**DOI:** 10.1101/801928

**Authors:** Chengchang Zhang, Meiling Lu, Lin Lin, Zhangjian Huang, Rongguang Zhang, Xuri Wu, Yijun Chen

## Abstract

As a vast repertoire of enzymes in nature, microbial cytochrome P450 monooxygenases require an activated form of flavin as a cofactor for the catalytic activity. Riboflavin is the precursor of FAD and FMN that serve as indispensable cofactors for flavoenzymes. In contrast to previous notion, here we describe the identification of an electron transfer process directly mediated by riboflavin for the N-dealkylation by microbial P450 monooxygenases. The electron relay from NADPH to riboflavin and then via activated oxygen to heme was proposed based on the combination of X-ray crystallography, molecular modeling and molecular dynamics simulation, site-directed mutagenesis and biochemical analysis of representative microbial P450 monooxygenases. This study provides new insights into the electron transfer mechanism in microbial P450 enzyme catalysis and likely in plants and mammals.

## Introduction

Cytochrome P450 monooxygenases ubiquitously occur in nature. P450s catalyze a variety of oxidative reactions, especially in the secondary metabolism in microorganisms and plants and drug metabolism in humans.^1, 2^ Recently, microbial P450s have become a major focus on the discovery of novel functionalities from the efforts of directed evolution and protein engineering.^3, 4^ Currently, there are two major electron transfer mechanisms for P450 enzymes.^5^ One is a system in which an NAD(P)H-dependent and FAD-containing reductase shuttles electrons to ferredoxin, which in turn transfers the electrons to P450 enzymes. Another is more general, which utilizes an NAD(P)H-dependent flavin mononucleotide (FMN) or flavin adenine dinucleotide (FAD)-containing reductase for the electron supply. Although most catalytic systems of P450s fit into above electron transfer pathways, several “unusual” electron transfer chains for different P450s have also been found, indicating that there still is much to be uncovered regarding the diversity of electron transfer for P450 enzymes.^5^ The most well-known and biotechnologically useful example for microbial P450s is the multi-domain fused and self-sufficient P450-BM3 from *Bacillus megaterium*.^6^ With regard to its electron transfer process, there have been quite a few studies. For example, a hinge-short-ended cytochrome P450 reductase ΔTGEE variant produces a stable complex of heme and hydroxyl radical and can support direct electron transfer from FMN to heme;^7^ the FMN domain of BM3 undergoes large conformational changes to bring FMN- and heme-domains close enough (8.8 Å) for facilitating the fast electron transfer in BM3.^8^ These studies have generated new knowledge on how the electrons are transferred during the catalysis by P450 enzymes. However, presently there is no general consensus on how flavin cofactors are participated in the reactions.

Typically, microbial P450s indispensably require a flavin cofactor, either FMN or FAD, for their electron transfer and catalytic activity regardless of being self-sufficient or the requirement of a redox partner.^9^ Regarding flavin cofactors, FAD and FMN are bio-synthesized from their precursor riboflavin (Vitamin B2, Figure 1), in which the isoalloxazine ring is a functional group to accept or transfer electrons.^10^ In flavoproteins, oxygen can be activated by the transfer of one electron from fully reduced flavin to O_2_.^11^ As previously known, the electrons originate from NAD(P)H that binds either transiently or permanently.^12^ Meanwhile, previous studies have indicated that riboflavin itself may possess the capability of accepting and transferring electrons from NAD(P)H. For instance, the heme domain of BM3 can be reduced by NADPH and FMN to yield the same product as did holo-BM3.^13^ Additionally, riboflavin could be used as an electron acceptor in an artificial flavoenzyme consisting of NADH and manganese porphyrin.^14^ Furthermore, riboflavin can donate electrons from NADH to a quinone in the NADH-driven sodium pump.^15^ Moreover, riboflavin can transfer electrons in modified CYP 2B4 and directly bind CYP 2B4.^16^ In the case of CYP 152, a microbial P450 enzyme, riboflavin could be activated by light-excitation and the electron donor ethylenediaminetetraacetate can serve as an electron source to complete the catalysis by CYP 152.^17^ Interestingly, in this artificial system, when riboflavin is free or covalently bound and either NAD(P)H or EDTA is used as the electron source, the peroxygenase activity is generally low, and the activity of CYP 152 remains significant even when high concentration of catalase is added into the reaction system, suggesting that the oxidative catalysis by CYP 152 is not related to possible peroxygenase activity. These collective evidences have implicated that riboflavin can directly participate in the electron transfer during P450 catalysis. However, how the electron transfer process is mediated by riboflavin in the complex catalysis remains elusive.

**Figure 1.**
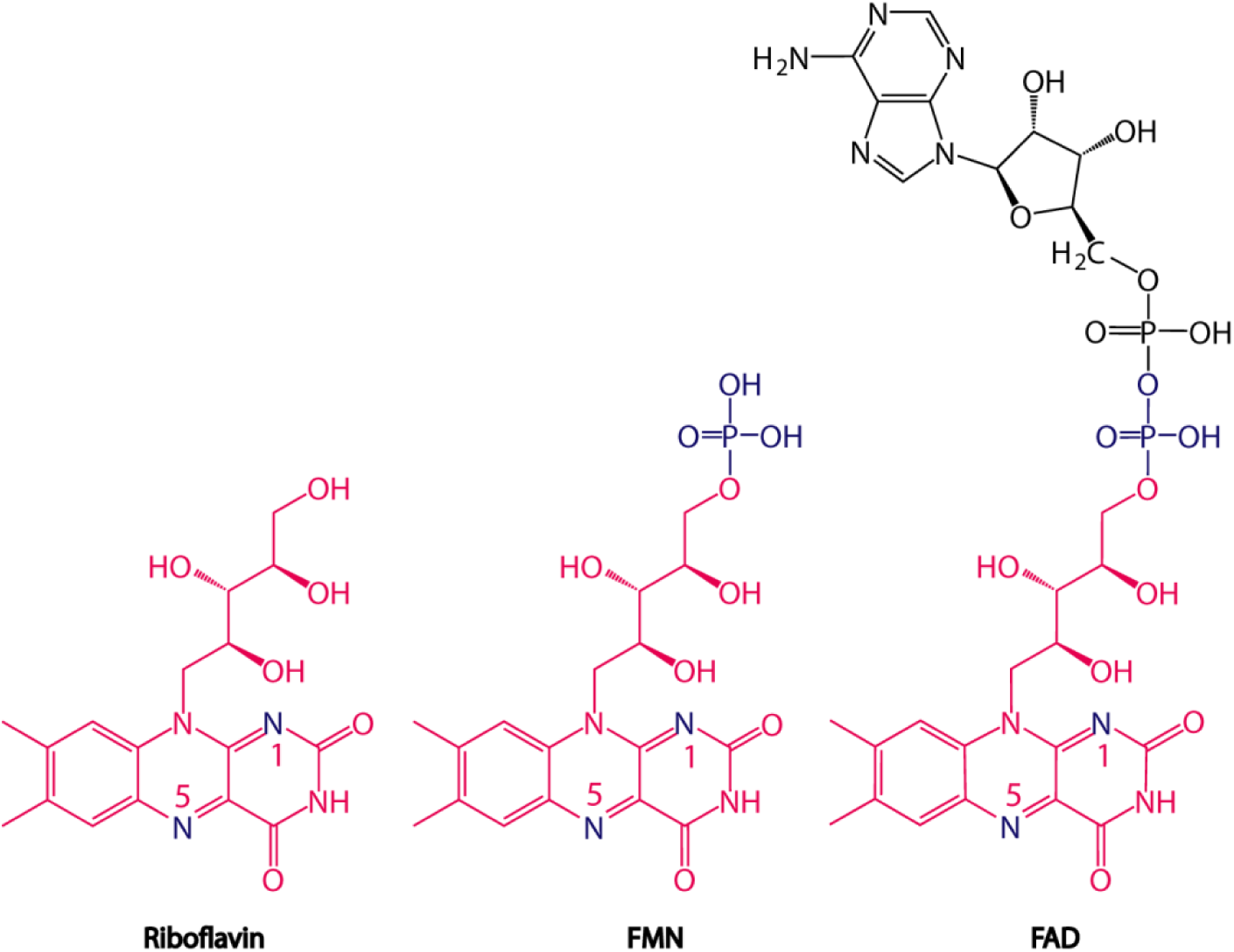
Structural comparison of riboflavin, FMN and FAD.

In the present study, given the valuable applications of N-dealkylation of P450s, especially in the area of drug metabolism,^18^ we chose this reaction based on the unusual finding as a model to uncover the roles of riboflavin in microbial P450s. After confirming riboflavin-mediated catalysis by representative microbial P450 monooxygenases, we were able to identify a new electron transfer pathway in addition to previously proposed catalytic mechanisms of P450s. This could enable our thorough understanding on various P450s and their catalytic mechanisms.

## Results

### Riboflavin directly mediated the N-dealkylation by microbial P450s

Given the occurrence of a large number of uncharacterized microbial P450 proteins in the databases, we first examined whether HmtS (GenBank: CBZ42153.1), a P450 homologous protein proposed to involve in the final step of the biosynthesis of himastatin,^19^ can utilize riboflavin for the natural reaction. Because the natural substrate of HmtS is difficult to obtain and impossible for chemical synthesis, we selected a series of simplified structural analogues to test the biaryl coupling. However, no biaryl products were found regardless of using riboflavin or ferredoxin/ferredoxin reductase (FDX/FDR) as a redox partner. Unexpectedly, N-dealkylation products were observed when riboflavin and NADPH were used as cofactors. Subsequently, after expression and purification of HmtS protein in a monomeric form with a molecular weight of 45 KDa (Figure S1), various substrates were screened for the dealkylation by HmtS in the presence of riboflavin and NADPH by LC-MS/MS (Table S1). Although eight compounds were identified to generate dealkylated products by HmtS, diphenhydramine was found to be the best substrate (Figure S2). In addition, the dealkylation reaction appeared both riboflavin- and NADPH-concentration dependent, and optimal pH and temperature for the reaction were determined to be pH7.4 and 34 °C respectively (Figure S3). After a preparative reaction with HmtS, diphenhydramine, riboflavin and NADPH, 50 mg N-desmethylated product was purified, and the chemical structure of the product was confirmed by NMR (Figure S4 and S5). Thus, diphenhydramine was used as a model substrate for subsequent examinations. Compared to FAD and FMN, riboflavin showed higher catalytic efficiency. Meanwhile, NADPH showed higher catalytic efficiency than NADH (Figure 2). To rule out the possibility of being a false positive, hemin, bovine serum albumin and bovine catalase were used to replace HmtS. Under the same reaction conditions, no product was observed in these non-catalytic components (Figure S6). These results strongly indicated that riboflavin indeed participates in the oxidative catalysis by HmtS.

**Figure 2.**
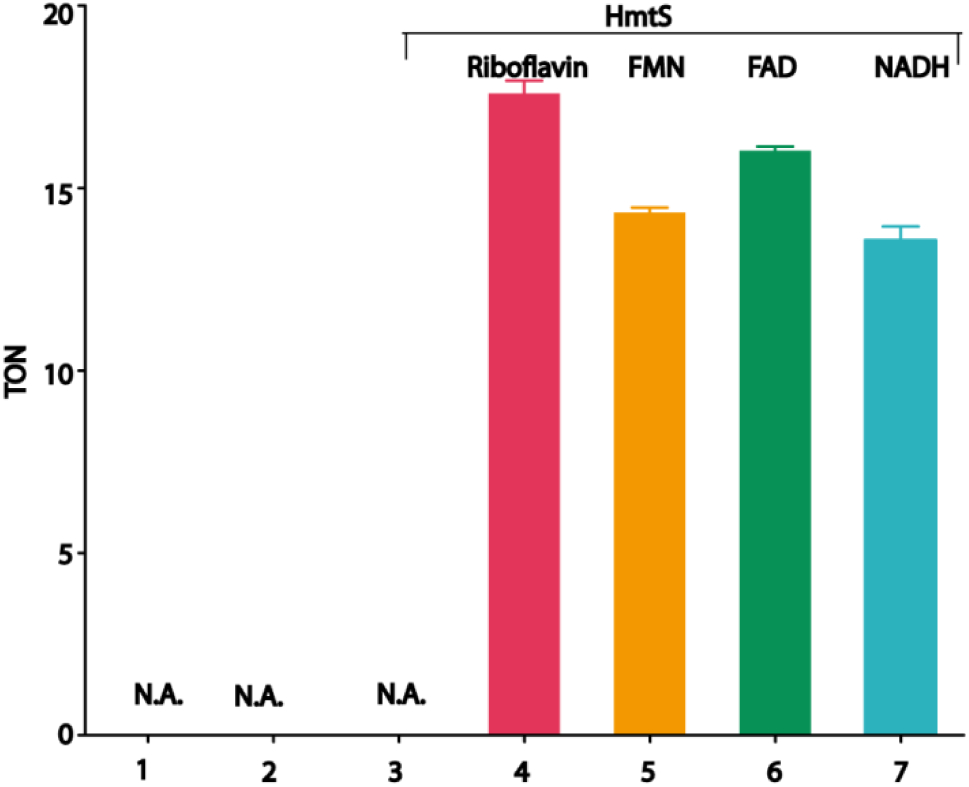
The N-dealkylation of diphenhydramine mediated by HmtS-riboflavin system. N. A. represents not active. Data are average values of 3 independent experiments with standard deviations. 1: NADPH; 2: NADPH + riboflavin; 3: enzyme + riboflavin; 4: enzyme + riboflavin + NADPH; 5: enzyme + FMN + NADPH; 6: enzyme + FAD + NADPH; 7: enzyme + riboflavin + NADH.

Next, four additional P450-like proteins that lack the reductase domain, including CYP107DY1 (UniProtKB: D5E3H2), ^20^ CYP106A2 (UniProtKB: Q06069), ^21^ HmtT (GenBank: CBZ42154.1) and HmtN (GenBank: CBZ42148.1),^19^ were further investigated to examine their capability of catalyzing the dealkylation. All four enzymes were active against diphenhydramine using riboflavin and NADPH as the cofactors and showed higher catalytic efficiency than HmtS (Figure S7a), suggesting that riboflavin can serve as the cofactor for a broad range of microbial P450-like monooxygenases that usually lack the reductase domain. To further confirm this finding and expand the scope of microbial P450s with riboflavin, the self-sufficient flavocytochrome P450 enzyme BM3^22, 23^ was evaluated by the comparison with its truncated mutant BMP lacking the reductase domain (BMR). As expected, the full-length BM3 containing BMR domain was able to catalyze the N-dealkylation of diphenhydramine while BMP was inactive. Intriguingly, BMP could catalyze the reactions in the presence of NADPH when riboflavin was added into the reaction mixtures, indicating that riboflavin could be used to functionally transfer electrons for the catalysis by BMP. Meanwhile, although FAD and FMN could also be functional for BMP, riboflavin showed higher efficiency on the enzyme catalysis (Figure S7b). These results confirmed again that riboflavin indeed participates in the electron transfer and enzyme catalysis. Therefore, the simplified catalytic system for microbial P450s consisted of enzyme, substrate, NADPH and riboflavin.

### Riboflavin effectively served as a redox partner for microbial P450s

It is well-known that the catalytic process by cytochromes P450 enzymes from various sources require a redox partner and NADPH reductase.^24^ Additionally, the binding of hydrogen peroxide to ferric heme is also described as the peroxide shunt pathway. Thus, we first compared riboflavin with classic redox partner FDR/FDX for the catalysis by HmtS. The initial rate with riboflavin for HmtS was lower than FDR/FDX, but the final conversion yield between these redox systems was equivalent (Figure S8). We next measured H_2_O_2_ formation during the catalysis by P450s with different redox partners. Significant amount of H_2_O_2_ was observed in FDR/FDX system (Figure S9a), whereas no measurable H_2_O_2_ was detected in the riboflavin system. Moreover, NADPH was significantly consumed in the riboflavin system compared to blank (Figure S9b). These observations suggested that the faster initial rate of FDR/FDX is likely due to the simultaneous generation of H_2_O_2_. Subsequently, to evaluate if the peroxide shunt pathway is involved in the catalysis by HmtS with riboflavin, exogeneous addition of H_2_O_2_ was performed to compare the catalytic efficiency. The catalytic efficiency of HmtS with riboflavin was generally higher than H_2_O_2_ (Figure S10). When catalase was added into the H_2_O_2_-mediated reactions, only small amount of the product was observed. On the other hand, the same amount of catalase did not significantly affect the catalysis with riboflavin (Figure S10). Collectively, the results indicated that the reaction mediated by riboflavin is not a result of producing H_2_O_2_, and suggested that riboflavin might mediate a new electron transfer pathway during the catalysis of HmtS.

### The enzymatic activity correlated to oxygen-binding and activation

To gain more information on the catalytic process of HmtS mediated by riboflavin, we aligned the amino acid sequence of HmtS with microbial P450 enzymes from different sources (Table S2). The sequence alignments indicated that HmtS contains a conserved oxygen-binding motif GXXT and a Leu residue at 340 (Figure S11), which is different from conserved Phe in the heme-binding motif of FXXGXXXCXG. This conserved Phe residue in P450s is responsible for modulating the reducing potential of heme through the interaction with conserved heme-biding residue Cys347.^25^ Therefore, HmtS-L340F mutant was constructed, expressed and purified. Both wild type (WT) and L340F mutant of HmtS were crystallized and their crystal structures (PDB: 5Z9I and 5Z9J) were solved at 2.0 Å and 2.2 Å respectively (Figure S12 and Table S3). Both structures adopt classic prism-shape architecture, consisting of 12 α helices and 14 β strands. The heme prosthetic groups are located in the typical binding pocket. Although overall structures of two proteins are very close (Figure S13), the difference is found in the pocket for O_2_ binding, in which the mutation from Leu to Phe results in a larger pocket due to the change of back-side region from heme. Meanwhile, the distance between heme and residue Thr241, the conserved site for O_2_ binding and activation,^26^ is obviously longer in HmtS-L340F than that in WT (Figure 3a). Compared to WT, L340F mutant showed approximately 3-fold increase of conversion (Figure S24), which may attribute the enlargement of O_2_ binding and activation site. Also, HmtS-L340F was more thermally stable (Figure S14). Additionally, spectroscopic analyses indicated that CO binding for HmtS-L340F mutant was much stronger than WT (Figure 3b) when dithionite was used as the reducing agent, suggesting that the activity increase by the mutant should be a result of the enlargement of O_2_ binding pocket^26^ for more rapid O_2_ activation. Together, the results suggested that the enlargement of oxygen-binding packet is critical to the catalytic activity with riboflavin.

**Figure 3.**
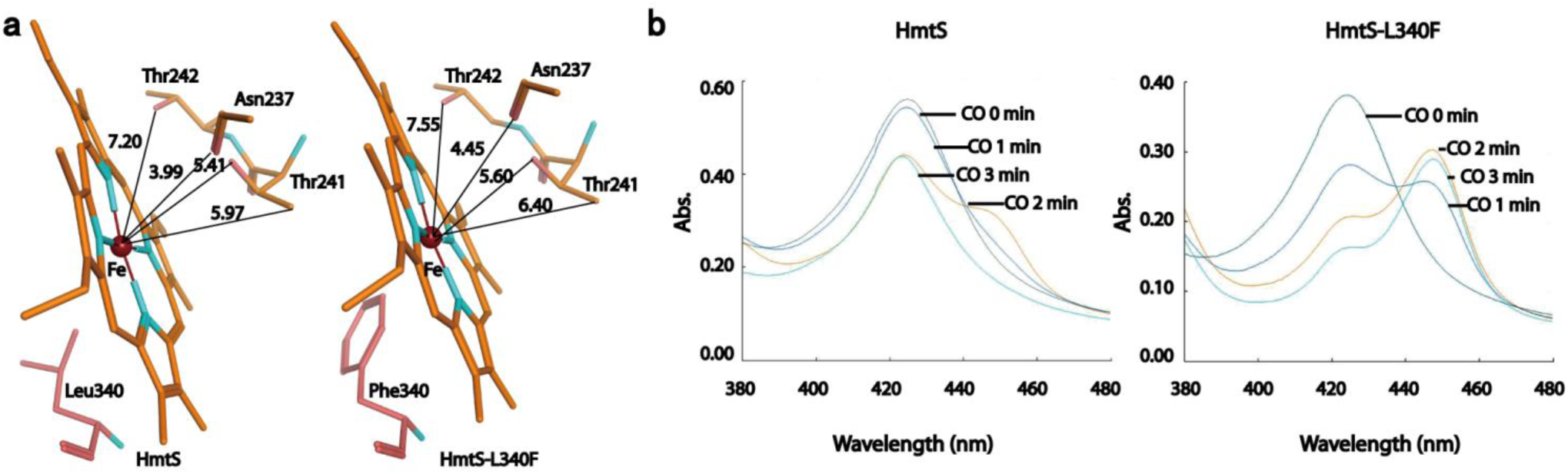
Comparison of HmtS and HmtS-L340F on the efficiency of utilizing O_2_. a) The enlargement of O_2_ binding pocket by L340F mutant. b) UV-Vis. spectroscopic differences on CO binding between HmtS and HmtS-L340F.

### Riboflavin mediated a new electron transfer pathway

To explore the electron transfer process by riboflavin, complexed structures of enzyme and cofactors were built by molecular modeling^27^ with the crystal structure of HmtS-L340F as a template due to its higher activity. In the modeled structures, HmtS-L340F can simultaneously accommodate and bind NADPH and riboflavin (Figure S15), suggesting that two cofactors can transfer electrons in the active pocket. Then, these models were used as initial conformations to conduct molecular dynamics (MD) simulation. According to literatures,^28^ 120-ns simulation for HmtS-L340F-riboflavin complex and 100-ns simulation for HmtS-L340F-riboflavin-NADPH complex were performed in Charm 3.6 field (Table S4). The changes in RMSD and RMSF indicated that both complexes can achieve equilibrium states in the initial stage by thermal energy (Figure S16). In the stimulation of HmtS-L340F-riboflavin complex, riboflavin can occupy the active pocket to form hydrogen bonds with residue Arg233 (Figure S17 and S18). However, there is a long distance between riboflavin and ferric ion of heme during the simulation process (Figure S19). Interestingly, when complexed with additional NADPH, the ribityl moiety in NADPH forms a new hydrogen bond with residue Asp74 (Figure S20). Subsequently, NADPH and riboflavin gradually adjust their positions in the active pocket in the initial stage of the simulation (Figure 4). Next, C4 of nicotinamide ring in NADPH moves closer to N1 of isoalloxazine ring of riboflavin to give a short distance of 4.81 Å between NADPH and riboflavin. Later, riboflavin gradually leaves from NADPH and moves to ferric ion of heme to yield an increased distance between NADPH and riboflavin (Figure 4b). As shown in Figure 4, the distance between riboflavin and heme is obviously decreased to 6.80 Å away from T240 residue after 100-ns simulation and remains unchanged. Consequently, N5 of isoalloxazine ring of riboflavin transfers electrons to ferric ion through O_2_ activation, which is stabilized and activated by Thr241 residue, to accomplish the N-dealkylation reaction. On the other hand, when the enzyme was complexed with the substrate in addition to riboflavin and NADPH, no stable confirmations were obtained during the 100 ns simulation and the substrate was not able to move close to heme (Figure S21), suggesting that the enzyme is unable to simultaneously bind the substrate and cofactors. This indicated that the dealkylation should be a sequential process. At the same time, NADPH exhibited two forms in the simulation of the electron transfer process (Table S5), suggesting that NADP^+^ may leave the active pocket after donating a pair of electrons to allow the entrance of the substrate for the completion of the catalytic cycle as proposed in Figure S22. Together, the MD simulation data provided evidence that NADPH directly interacts riboflavin for transferring electrons to heme, and then oxidizes the substrate for the catalysis.

**Figure 4.**
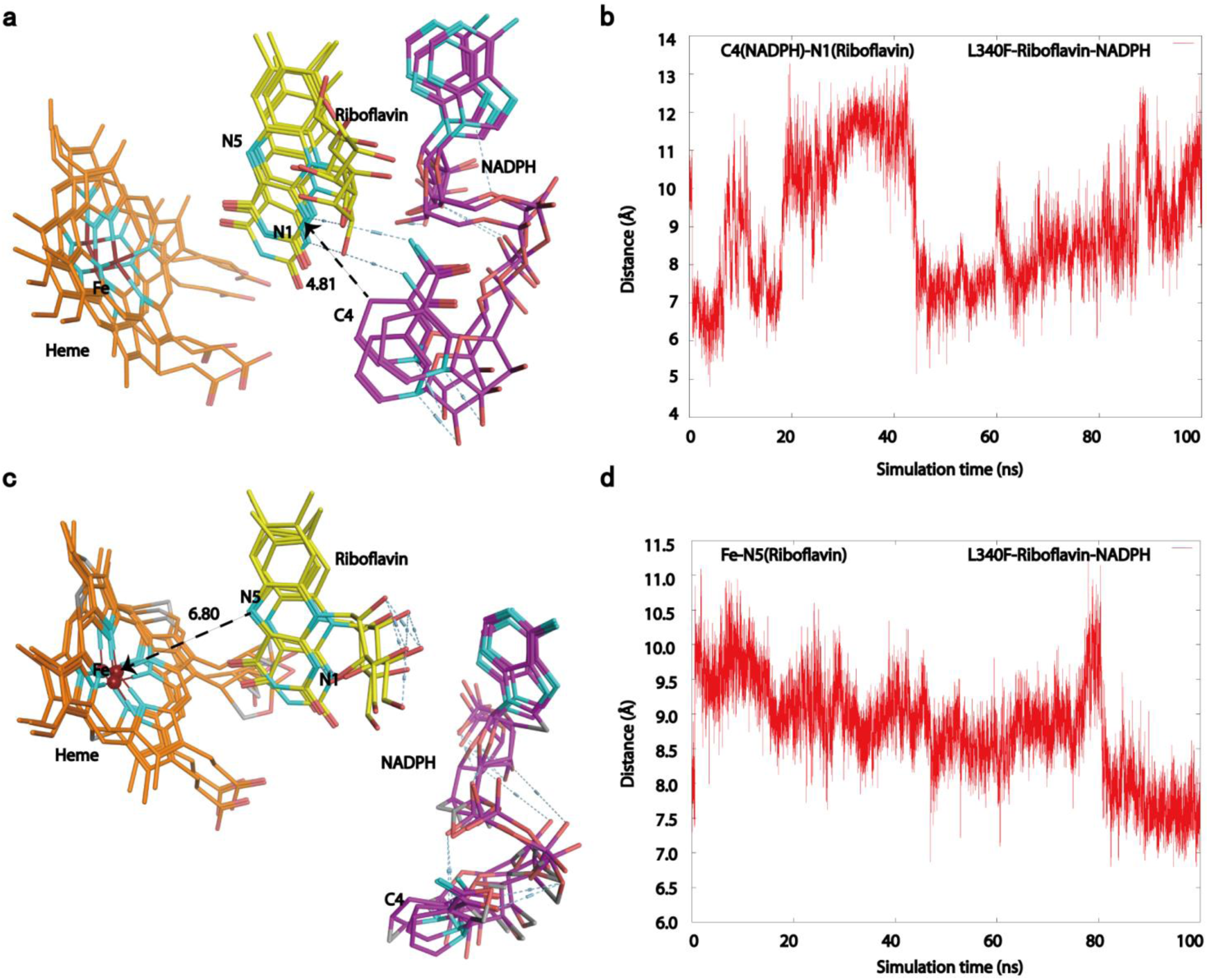
MD simulation for the complexed models and the electron transfer process mediated by HmtS-L340F. a) Serial spatial positions between NADPH and riboflavin in HmtS-L340F-riboflavin-NADPH complex. b) Distance fluctuations of C4 (NADPH) to N1 (riboflavin) in HmtS-L340F-riboflavin-NADPH complex. c) Serial spatial positions between riboflavin and heme in HmtS-L340F-riboflavin-NADPH complex. d) Distance fluctuations of N5 (riboflavin) to Fe center in HmtS-L340F-riboflavin-NADPH complex. Ribbon diagram of NADPH (violet) and riboflavin (yellow) is shown in the models.

### Biochemical analyses confirmed the electron transfer process

After excluding the possibility of dimerization (Figure S1 and S23) and the existence of covalently bound flavins in the enzyme (Figure S24), to verify the proposed electron transfer process, we individually tested whether the substrate or cofactors can directly bind the enzyme measured by Microscale Thermophoresis Assay (MST). Although the binding affinity between HmtS and substrate or NADPH was relatively weak, the affinity between HmtS and riboflavin was strong (Figure S25), which is similar to pervious report.^29^ Meanwhile, the binding affinity of riboflavin was also slightly stronger than FAD or FMN, suggesting a possible role of steric hindrance (Figure S25).

Based on sequence alignments (Figure S11), molecular modeling (Figure S15) and MD simulation (Figure 4), among sixteen residues in HmtS for potential involvement in the binding and catalysis as listed in Table S6, four important residues, including Cys347 for heme binding, Thr241 for O_2_ binding and activation, Arg233 putatively for riboflavin binding and Asp74 putatively for NADPH binding, were identified and individually mutated to Ala to examine their influences to the activity. As anticipated, in addition to conserved residues of Cys347 and Thr241, both Arg233 and Asp74 could remarkably affect the activity of HmtS-L340F from the mutagenesis. In the complexed model consisting of enzyme, NADPH and riboflavin, Arg233 is the site to bind riboflavin (Figure S17 and S18), and Asp74, similar to the situation of Asp632 in NADPH-cytochrome P450 oxidoreductase,^30^ is responsible for the binding with NADPH (Figure S20). Not surprisingly, the significant reduction of activity from the mutations of Arg233 and Asp74 was in accordance with the results of MD simulation. Subsequently, double mutation of R233A and D74A indeed resulted in the complete loss of activity (Figure S26). To examine whether superoxide anion is involved in the oxidative catalysis, the reaction was completely abolished when 1 U/ml of superoxide dismutase (SOD) from bovine erythrocytes was added into the reaction system consisting of HmtS-L340F, diphenhydramine, riboflavin and NADPH (Figure S26), conforming the generation of superoxide anion in the catalytic process. The collective evidences not only demonstrated the importance of Arg233 and Asp74 residues as the binding sites for the cofactors, but also probed the involvement of superoxide anion, which strongly supports the proposed mechanism of electron transfer process mediated by riboflavin as illustrated in Figure 5.

**Figure 5.**
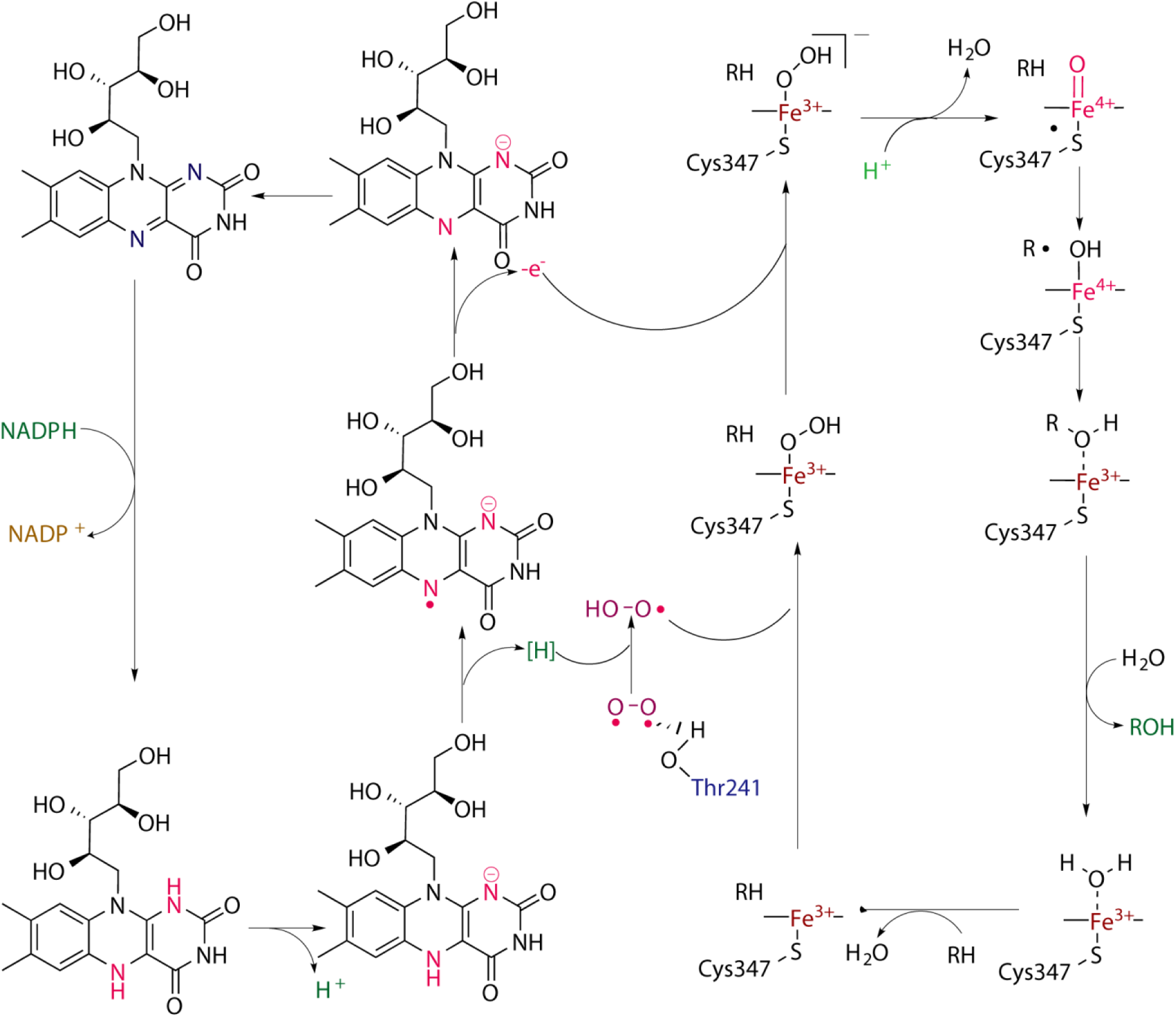
Proposed mechanism for the electron transfer by riboflavin in microbial P450 monooxygenases.

## Discussion and conclusions

Cytochrome P450 monooxygenases are heme-containing enzymes that use molecular oxygen and NAD(P)H for oxidative insertion of one oxygen atom into an organic substrate. Typically, P450 enzymes exhibit poor stability, low turnover number and require electron transport protein(s) for their catalysis. Among different classes of P450s, class I enzymes encompass most of known bacterial P450s in addition to a number of eukaryotic mitochondrial P450s. This class of enzymes usually possess a characteristic of two P450 components, a FAD-dependent ferredoxin reductase and a ferredoxin containing an iron-sulfur cluster and require NAD(P)H as a source of electrons.^5^ Previously, HmtS was proposed to involve in the biosynthesis of himastatin,^19^ presumably responsible for the biaryl coupling. However, its actual role has not been experimentally confirmed. Given this interesting reaction, we have attempted to verify the biaryl coupling *in vitro*, but the efforts have been unsuccessful. On the other hand, when we screened various substrates for N-dealkylation, HmtS exhibited notable activity. Therefore, based on the structural features that, like widely existed microbial P450s, lack the reductase domain, we selected this microbial P450 as a model enzyme to probe its cofactor requirement, particularly whether riboflavin could substitute classic redox partner for the catalysis based on previous observations.^12-16^ Surprisingly, HmtS indeed could directly utilize riboflavin and NADPH to accomplish the N-delakylation, showing higher efficiency than FAD and FMN. With regard to the natural substrate and possible involvement in the biaryl coupling by HmtS in the organism, we are uncertain at this point, which requires further investigation. However, a thorough comparison revealed that this dealkylation reaction is mediated by a new electron transfer pathway, which is not the same as any reported mechanisms for P450s.^31^ Subsequently, based on sequence alignments with other microbial P450s, we found that residue Leu340 of HmtS is different from other enzymes. Thus, HmtS-L340F mutant was constructed to show higher catalytic efficiency. To elucidate possible mechanism on the electron transfer, both HmtS and HmtS-L340F were crystallized and their crystal structures were solved. Although the complexed structures of the enzyme with riboflavin and NADPH were unable to obtain after numerous attempts due likely to the relatively lower binding affinity, we were able to identify the electron transfer process through MD simulations, and a new catalytic mechanism was then proposed, which may possess a variety of implications for various P450s. Of note, same as most other P450s, the catalytic efficiency of HmtS was relatively low, which might be a result of stability issue and the complex electron transfer process required by multi-components.

As one of the common oxidative reactions, N-dealkylation can be catalyzed by various oxygenases. However, regioselectivity is still an issue for many enzymes on their biocatalytic applications. By contrast, the present regioselective dealkylation by HmtS, riboflavin and NADPH could be utilized to prepare specific amines to serve as versatile building blocks for the synthesis of various pharmaceuticals and bioactive compounds.^32^ In addition, this system could replace microsomal preparations and mammalian P450s for the generation of relatively large quantity of drug metabolites in a short period of time for preclinical evaluation of drug metabolism and toxicity,^33^ which would be helpful to speed up the process of drug discovery.

The studies on cytochrome P450 monooxygenases have increasingly been a spotlight,^34, 35^ especially the exploration and discovery of the catalysis of novel reactions by protein engineering. A recent report on the α-Amino C-H fluoroalkylation by engineered P450 enzymes^36^ is just one of the excellent examples. Meanwhile, regardless of *de novo* design or rational engineering of enzymes, the catalytic mechanisms are a perquisite. Thus, we believe that our present finding can also provide a starting point for the design and engineering of different P450 enzymes, and we anticipate that much more novel enzymes and reactions are yet to come.

In summary, we have demonstrated that riboflavin directly participates in the oxidative catalysis by microbial cytochrome P450 monooxygenases. Based on the collective evidences, we have proposed a new electron transfer pathway for the N-demethylation reaction by microbial P450 enzyme, NADPH and riboflavin, which might be generally applicable to numerous P450-like monooxygenases that lack the reductase domain. The present work has established a foundation to better understand cofactor requirement, catalytic mechanism and substrate scope for numerous P450-like enzymes in nature.

## Experimental section

### Construction of plasmids and site-directed mutagenesis

PCR products containing *fdr, fdx, cyp106A2, cyp107DY1, cyp102A1, BMP, hmtT, hmtN* and *hmtS* were each amplified and cloned into the *Nde I*/*Xho I* or *Nde I*/H*in*d III sites of pET28a(+) or pET22b(+) (ClonExpress II One Step Cloning Kit Vazyme Biotech Co. Ltd, China) to produce the recombinant constructs pFDR-ET28a, pFDX-ET28a, pBM3-ET28a, pBMP-ET28a, pCYP106A2-ET28a, pCYP107DY1-ET28a, pHmtT-ET28a, pHmtN-ET28a, pHmtS-ET22b and pHmtS-ET28a for the expression of FDR, FDX, CYP106A2, CYP107DY1, P450 BM3, BMP, HmtT, HmtN and HmtS, respectively. Mutagenesis was performed with pHmtS-ET22b as a template using a QuickChange site-directed mutagenesis kit (Stratagene, La Jolla, USA) according to the manufacturer’s protocol. Superoxide dismutase (SOD) from bovine erythrocytes was purchased from Aladdin (Shanghai, China).

### Enzyme activity assay

The reactions for the screening of various substrates were performed for 4 h using 4 μM enzyme, 3 mM NADPH, 150 μM substrate and 4 μM riboflavin in PBS buffer (137 mM NaCl, 2.7 mM KCl, 10 mM Na_2_HPO_4_, 2 mM KH_2_PO_4_, pH 7.4). Most substrates were dissolved in the buffer, and some with addition of 0.4% DMSO or methanol. The reaction mixtures contained 4 μM purified enzyme, 3 mM NADPH, 4 μM riboflavin and 400 μM substrate in PBS buffer (pH7.4) to a final volume of 200 μL, and aliquoted in triplicate into a 48-well plate for 4-h reactions. The catalytic efficiency of all P450-riboflavin systems was expressed by turnover number (TON). To avoid possible interference from light, the plates containing reaction mixtures were covered and wrapped in foil. After shaking the plates at 250 rpm, the reactions were quenched by addition of methanol (400 μL) at −20 °C. This mixture was then transferred to a microcentrifuge tube and centrifuged at 12,000 × g for 10 min. The supernatant was subjected to HPLC analysis. Conversion and TON were calculated based on the peak areas of product and substrate with standard calibration curves.

### Microscale thermophoresis (MST) assay

Purified enzymes were labeled with primary-amine coupling of NT647 dye according to the manufacturers’ protocol and purified from free dye with Superdex 25 resin in the buffer (50 mM HEPES, pH 7.5, 100 mM NaCl) at 25 °C. Titration series were made with 1:1 dilution of ligands while maintaining a constant 50 nM NT-647 labeled enzymes. The initial concentrations of diphenhydramine, NADPH, riboflavin, FAD and FMN were 2 mM, 1 mM, 250 μM, 250 μM and 250 μM, respectively. The sample was loaded into the NanoTemper glass capillaries and microthermophoresis carried out using 20% LED power and 80% MST. K_d_ values were calculated using the mass action equation *via* the NanoTemper software from duplicate reads of triplicate experiments.

### UV-Visible spectroscopic analysis

CO binding spectra were recorded as described previously.^37^ UV-Visible spectra of HmtS and HmtS-L340F were recorded from 300 to 550 nm. The ferric heme was reduced by adding freshly prepared saturated solution of dithionite (Na_2_S_2_O_4_) to the protein solution, then was bubbled softly by CO gas, and the spectrum of reduced enzyme was then recorded.

### Protein crystallization and structural determination

After the screening of crystallization conditions, HmtS crystals were obtained using a reservoir solution of 22% (w/v) polyethylene glycol (PEG) 3350, 0.1 M ammonium acetate, 0.1 M NDSB-256 and 0.1 M Bis-Tris (pH 5.5) and grown at 20 °C by the sitting-drop vapor diffusion method, in which 1 μL of 6 mg/ml protein solution in 0.1 M NaCl, 1 mM DTT, 20 mM Tris-HCl buffer (pH 7.4) was mixed with an equal volume of the reservoir solution. For the preparation of HmtS-L340F crystals, the reservoir solution consisted of 20% (w/v) PEG3350, 0.15 M ammonium sulfate and 0.1 M Bis-Tris (pH 6.5). Other solutions and experimental methods were same as those used for HmtS crystallization.

Crystals were protected by using 25% glycerol as the cryoprotectant. The diffraction data were collected at 19 U Beamline of Shanghai Synchrotron Radiation Facility (SSRF), and were processed by HKL2000 package.^38^ Molecular replacement was performed by Phaser in CCP4^39^ to solve the structure of HmtS-L340F using the crystal structure of HmtT (PDB: 4GGV) as the searching model. The structure of HmtS was solved by molecular replacement using the mutant structure as a searching model. Refinements were carried out using Phenix. refine.^40^ Coot was used for model buildings and modifications.^41^ For details, see Supplementary Table S3. Crystal structures of HmtS and HmtS-L340F were overlapped and the O_2_ binding pocket of HmtS and HmtS-L340F was determined from the distances of heme-N237, heme-T242, heme-T241 using MOE2014.

### Molecular dynamics simulation

To verify the modeled structures and probe the catalytic process, molecular dynamics (MD) simulation for the complexes of HmtS-L340F-riboflavin and HmtS-L340F-riboflavin-NADPH were started from initial conformations. The MD simulation was performed with GROMACS package. The Charmm3.6 field was used for generating the topology and coordinate files for each system. All small molecule force fields including riboflavin, NADPH and heme in HmtS-L340F were generated by CGENFF force field file, and then converted into the format supported by GROMACS package. Each structure was immersed in a regular dodecahedron box of TIP3P waters, resulting in the addition of 12,400 solvent molecules on average. Each system was neutralized through the addition of counter ions and energy minimization. To avoid the dissolution of small molecules in the equilibrium process, protein, riboflavin and NADPH were classified as one group, and solvent and ions as another group. Subsequently, the systems were gradually heated from 0 K to 300 K for 5 ns of density equilibration in the NVT ensemble with a 2-fs time step, then the systems were equilibrated in NPT ensemble at a target temperature of 300 K and a target pressure of 1.0 bar with a 2-fs time step under periodic boundary conditions. Totally, HmtS-L340F-riboflavin complex was simulated for 120 ns, and HmtS-L340F-riboflavin-NADPH complex was simulated for 100 ns. The results were analyzed, focusing on the changes of the distance between key atoms in the simulated period and the conformational changes of small molecules and HmtS-L340F.

## Author contributions

Y. C. and X. W. conceived and supervised the research. C. Z. and M. L. designed and performed the experiments and analyzed the data, L.L. and R. Z. solved the crystal structures. Z. H. analyzed data. C. and Y.C. wrote the manuscript. ^**‡**^C. Z and M. L contributed to the work equally.

## Additional information

Experimental procedures, characterization of small molecule compounds, UV-Visible spectroscopy, additional data of HPLC, LC-MS/MS, structural data of the enzymes and MD simulation are included in the supporting information.

## Acknowledgements

This work is supported by the National Key R&D Program of China (2018YFA0902000), the “111 Project” from the Ministry of Education of China and the State Administration of Foreign Expert Affairs of China (No. 111-2-07), “Double First-Class” University project (CPU2018GY36) and National Science Foundation of China (No: 21778076). We are grateful to the staff of beamline BL19U at SSRF, Shang-hai, China, for the assistance on X-ray crystallography.

