## supplemental file for "Riboflavin Directly Mediates the Dealkylation by Microbial Cytochrome P450 Monooxygeneses"

### 1. Supplementary Methods

#### Protein expression and purification

*E. coli* BL21 (DE3) cells were transformed with the expression plasmids. A single colony transformant was selected to inoculate a 5 ml culture of LB medium supplemented with 50 μg/ml kanamycin. The culture was grown at 37 ℃ for 12 h and used to inoculate 1 L of LB medium containing 50 μg/ml kanamycin. Cells were grown at 37 ℃ and 220 rpm to $OD_{600}$ = 0.6-1.5, and then IPTG was added to a final concentration of 0.1-0.5 mM. After additional 5-8 h of incubation at 16 ℃, the cells were harvested by centrifugation at 4,000 × g for 15 min at 4 ℃. The pellets were used directly for protein purification or stored at -80 ℃ before use.

The cell pellets collected by centrifugation were re-suspended in lysis buffer (25 mM Tris-HCl, 20 mM imidazole, 50 mM NaCl, pH 7.5), and were disrupted at 4 ℃. Then, cell debris was removed via centrifugation at 21,000 × g for 30 min at 4 ℃. The resulting supernatant was incubated with Ni-NTA resin pre-equilibrated with the lysis buffer, and then subjected to affinity purification on a column. The desired elution fractions with different concentrations of imidozole were combined and concentrated using an Amicon Ultra-15 Centrifugal Filter Unit (Millipore, USA). The concentrated protein solution was desalted using Sephadex$^{TM}$ G25 column (GE Healthcare, USA). For the enzymes used for the catalytic reactions and all biochemical and biophysical analyses, the affinity chromatography and the ultrafiltration with an Amicon Ultra-15 Centrifugal Filter Unit (Millipore, USA) were performed for 3 times to obtain electrophoresitically pure proteins (Figure S1). Recombinant HmtS and HmtS-L340F, used for crystallization, were further purified by gel filtration chromatography (HiLoad 16/600

Superdex 200 pg column, GE Healthcare) after His-tag cleavage by thrombin. The purity of the protein samples was examined by 10% SDS-PAGE and 10% native PAGE. The final proteins used for crystallization were concentrated to 10 mg/ml and flash-frozen by liquid nitrogen and stored at -80 ℃.

**Analytical HPLC method**

All HPLC methods were performed with various gradients of mobile phase A (0.1% TFA in water) and mobile phase B (0.1% TFA in acetonitrile) at 28 ℃ and 10 μl of sample injection volume with UV-DAD detection. The analytical procedures were as follows.

General HPLC analyses for substrate screening:

| Time (min) | Mobile phase A | Mobile phase B |
|---|---|---|
| 0.00 | 95.0% | 5.0% |
| 5.00 | 95.0% | 5.0% |
| 25.00 | 15.0% | 85.0% |

HPLC analyses for diphenhydramine and its product:

| Time (min) | Mobile phase A | Mobile phase B |
|---|---|---|
| 0.00 | 95.0% | 5.0% |
| 5.00 | 95.0% | 5.0% |
| 6.00 | 65.0% | 35.0% |
| 18.00 | 65.0% | 35.0% |

**LC-MS/MS method**

The mobile phase used for HPLC separation consisted of 0.1% formic acid in water (A) and 0.1% formic acid in acetonitrile (B). The HPLC conditions were the same as that for substrate screening. The MS/MS parameters for the product identification were set as follows.

Parameters for MS/MS detection:

| Parameter | Value |
|---|---|
| MS Min Range | 100 |
| MS Max Range | 1700 |
| MS Scan Rate | 4 |
| MS/MS Min Range | 100 |
| MS/MS Max Range | 1700 |
| MS/MS Scan Rate | 2 |
| Fixed Collision Energy | 30 |
| Ion Polarity | Positive/Negative |
| Ion Source | Dual ESI |

**Preparation of N-dealkylation product from diphenhydramine**

The reaction was conducted with 680 μM diphenhydramine, 5 μM enzyme, 5 mM NADPH and 5 μM riboflavin in 20 mM PBS buffer (pH 7.4) to a final volume of 50 ml for 8 h. The reaction mixture was extracted by equal volume of ethyl acetate twice to collect crude product after concentration under reduced pressure. The crude product was further purified by preparative HPLC (35% acetonitrile as mobile phase at a flow rate of 10 ml/min). Final product (50 mg) was obtained after freeze-dry with a purity of 98%.

**Determination of native molecular weight of HmtS**

Sephacryl S-200 HR column was employed to determine the molecular weight of purified HmtS. The calibration curve of proteins with standard molecular weight was obtained using gel filtration calibration kits (GE Healthcare, USA) according to the manufacture's protocol.

**Measurement of thermal stability**

The detection conditions were typically as follows: heating range 35 ℃ - 95 ℃ for 3 min at protein concentration of 0.08 mg/ml. In this study, fluorescent dye labeled HmtS-L340F was added as a control to ensure the accuracy of the detection.

**Determination of $H_2O_2$ concentration**

The assay was carried out in PBS buffer pH 7.4 (final volume 190 μl), and contained 118 μl of incubated reaction mixture as indicated in Figure S9a. Final concentrations of other components were 125 μM phenol, 1.25 mM 4-aminoantipyrine, and 0.1 mg/ml horseradish peroxidase. The absorbance at 510 nm (λmax of the quinoneimine product) was recorded and the concentration of $H_2O_2$ in each reaction mixture was calculated from a calibration curve using serial concentrations of $H_2O_2$. To verify if the color shift observed in the reaction attributed $H_2O_2$, samples containing catalase (20 U/ml) were assayed as controls.

### 2. Supplementary Tables

**Table S1.** Screening of different substrates for HmtS-L340F with riboflavin

| Substrate | Exact mass | Observed mass (m/z) | Major ions (m/z) | Conversion (%) |
|---|---|---|---|---|
| **1** | 241.15 | 242.15 | 228.14, 152.06 | 68 |
| **2** | 241.15 | 242.15 | 163.13, 134.09 | 63 |
| **3** | 246.10 | 247.10 | 230.07, 167.07 | 10 |
| **3** | 260.11 | 261.11 | 230.07, 167.07 | 45 |
| **4** | 299.15 | 300.16 | 209.10, 133.06, 91.05 | 18 |
| **5** | 160.10 | 161.11 | 117.06, 77.04 | 5 |
| **6** | 165.12 | 166.12 | 107.05, 58.06 | 8 |
| **7** | 141.03 | 142.04 | 125.02, 99.00 | 16 |
| **8** | 141.03 | 142.04 | 125.02, 99.00 | 12 |

**Table S2.** List of P450 enzymes and their Uniprot entry names.

| P450 enzymes | Uniprot entry name | Source |
|---|---|---|
| HmtT | D9WMR2 | *Streptomyces himastatinicus* ATCC 53653 |
| KtzM | A8CF80 | *Kutzneria* sp. 744 |
| HmtN | D9WMQ6 | *Streptomyces himastatinicus* ATCC 53653 |
| HmtS | G0LWB2 | *Streptomyces himastatinicus* ATCC 53653 |
| CYP107DY1 | D5E3H2 | *Bacillus megaterium* ATCC 12872 |
| CYP106A2 | Q06069 | *Bacillus megaterium* Taxonomic (NCBI) 1404 |
| CYP101 | P00183 | *Pseudomonas putida* Taxonomic (NCBI) 303 |
| CYP102A1 | P14779 | *Bacillus megaterium* ATCC 14581 |

**Table S3.** Data collection and refinement statistics of HmtS (PDB: 5Z9I) and HmtS-L340Fmutant (PDB: 5Z9J).

| Data collection | | |
|---|---|---|
| Dataset | HmtS | HmtS-L340F |
| Space group | P1 | $P2_12_12_1$ |
| Unit cell (a, b, c, Å) | 57.0, 78.5, 97.7 | 57.9, 72.4, 98.4 |
| Unit cell (α, β, γ,°) | 78.3, 72.6, 69.5 | 90, 90, 90 |
| Wavelength (Å) | 0.9790 | 0.9778 |
| Resolution range (Å) | 50.00-2.00 (2.03-2.00) | 50.00-2.20 (2.33-2.20) |
| No. of unique reflections | 92892 (4120) | 40569 (6635) |
| Redundancy | 3.7 (3.1) | 3.4 (3.2) |
| I/σ | 15.1 (1.9) | 13.2 (2.9) |
| Completeness (%) | 90.1 (80.1) | 99.1 (97.1) |
| Rmerge (%) [a] | 7.8 (38.6) | 6.5 (44.0) |
| Structure refinement | | |
| Resolution, Å | 46.33-2.00 | 45.28-2.20 |
| Rwork[b]/Rfree[c] (%) | 22.20/26.16 | 18.31/23.36 |
| Rmsd bonds/angles (Å/ °) | 0.013/1.478 | 0.009/1.471 |
| Average B factor ($Å^2$) | 43.0 | 46.0 |
| No. of protein atoms | 11727 | 2911 |
| No. of HEM molecules | 172 | 43 |
| No. of solvent molecules | 565 | 114 |
| No. of reflections | 87214 | 40568 |
| Ramachandran plot (%) | | |
| Favored regions | 97.3 | 95.6 |
| Allowed regions | 2.7 | 4.4 |
| Outliers | 0 | 0 |

Numbers in parentheses represent the value for the highest resolution shell.

[a] Rmerge = ∑|Ii - Im|/∑Ii, where Ii is the intensity of the measured reflection and Im is the mean intensity of all symmetry related reflections.

[b] Rcryst = Σ||Fobs| - |Fcalc||/Σ|Fobs|, where Fobs and Fcalc are observed and calculated structure factors.

[c] Rfree = ΣT||Fobs| - |Fcalc||/ΣT|Fobs|, where T is a test data set of about 5% of the total reflections randomly chosen and set aside prior to refinement.

**Table S4.** Force field parameters for heme.

| Index | Name | Coordinate x | Coordinate y | Coordinate z | Type | | | Partial charges |
|---|---|---|---|---|---|---|---|---|
| 1 | FE | -1.211 | -1.295 | 1.717 | FE | 1 | HEM | 0.24 |
| 2 | N1 | -1.267 | -1.344 | 1.897 | NPH | 1 | HEM | -0.18 |
| 3 | N2 | -1.147 | -1.113 | 1.789 | NPH | 1 | HEM | -0.18 |
| 4 | N3 | -1.172 | -1.216 | 1.531 | NPH | 1 | HEM | -0.18 |
| 5 | N4 | -1.285 | -1.447 | 1.643 | NPH | 1 | HEM | -0.18 |
| 6 | C1 | -1.312 | -1.465 | 1.930 | CPA | 1 | HEM | 0.12 |
| 7 | C2 | -1.333 | -1.472 | 2.066 | CPB | 1 | HEM | -0.06 |
| 8 | C3 | -1.298 | -1.354 | 2.114 | CPB | 1 | HEM | -0.06 |
| 9 | C4 | -1.257 | -1.275 | 2.009 | CPA | 1 | HEM | 0.12 |
| 10 | C5 | -1.157 | -1.076 | 1.914 | CPA | 1 | HEM | 0.12 |
| 11 | C6 | -1.102 | -0.951 | 1.929 | CPB | 1 | HEM | -0.06 |
| 12 | C7 | -1.060 | -0.913 | 1.809 | CPB | 1 | HEM | -0.06 |
| 13 | C8 | -1.089 | -1.020 | 1.719 | CPA | 1 | HEM | 0.12 |
| 14 | C9 | -1.103 | -1.109 | 1.498 | CPA | 1 | HEM | 0.12 |
| 15 | C10 | -1.085 | -1.107 | 1.363 | CPB | 1 | HEM | -0.06 |
| 16 | C11 | -1.145 | -1.215 | 1.314 | CPB | 1 | HEM | -0.06 |
| 17 | C12 | -1.199 | -1.283 | 1.420 | CPA | 1 | HEM | 0.12 |
| 18 | C13 | -1.301 | -1.475 | 1.516 | CPA | 1 | HEM | 0.12 |
| 19 | C14 | -1.363 | -1.602 | 1.499 | CPB | 1 | HEM | -0.06 |
| 20 | C15 | -1.384 | -1.648 | 1.619 | CPB | 1 | HEM | -0.06 |
| 21 | C16 | -1.334 | -1.549 | 1.710 | CPA | 1 | HEM | 0.12 |
| 22 | C17 | -1.343 | -1.565 | 1.845 | CPM | 1 | HEM | -0.1 |
| 23 | C18 | -1.372 | -1.653 | 1.882 | HA | 1 | HEM | 0.1 |
| 24 | C19 | -1.211 | -1.148 | 2.021 | CPM | 1 | HEM | -0.1 |
| 25 | H1 | -1.216 | -1.103 | 2.110 | HA | 1 | HEM | 0.1 |
| 26 | C20 | -1.061 | -1.015 | 1.586 | CPM | 1 | HEM | -0.1 |
| 27 | H2 | -1.006 | -0.939 | 1.552 | HA | 1 | HEM | 0.1 |
| 28 | C21 | -1.264 | -1.401 | 1.408 | CPM | 1 | HEM | -0.1 |
| 29 | H3 | -1.286 | -1.435 | 1.317 | HA | 1 | HEM | 0.1 |
| 30 | C22 | -1.301 | -1.309 | 2.255 | CT3 | 1 | HEM | -0.27 |

| | | | | | | | | |
|---|---|---|---|---|---|---|---|---|
| 31 | C23 | -1.269 | -1.215 | 2.261 | HA3 | 1 | HEM | 0.09 |
| 32 | H4 | -1.394 | -1.315 | 2.290 | HA3 | 1 | HEM | 0.09 |
| 33 | H5 | -1.241 | -1.368 | 2.310 | HA3 | 1 | HEM | 0.09 |
| 34 | C24 | -1.385 | -1.592 | 2.141 | CT2 | 1 | HEM | -0.18 |
| 35 | H6 | -1.334 | -1.598 | 2.227 | HA2 | 1 | HEM | 0.09 |
| 36 | H7 | -1.367 | -1.673 | 2.086 | HA2 | 1 | HEM | 0.09 |
| 37 | C25 | -1.533 | -1.591 | 2.174 | CT2 | 1 | HEM | -0.28 |
| 38 | H8 | -1.585 | -1.593 | 2.089 | HA2 | 1 | HEM | 0.09 |
| 39 | H9 | -1.554 | -1.506 | 2.224 | HA2 | 1 | HEM | 0.09 |
| 40 | C26 | -1.578 | -1.707 | 2.259 | CC | 1 | HEM | 0.62 |
| 41 | O1 | -1.694 | -1.731 | 2.271 | OC | 1 | HEM | -0.76 |
| 42 | O2 | -1.501 | -1.777 | 2.318 | OC | 1 | HEM | -0.76 |
| 43 | C27 | -1.095 | -0.874 | 2.056 | CT3 | 1 | HEM | -0.27 |
| 44 | H10 | -1.050 | -0.786 | 2.039 | HA3 | 1 | HEM | 0.09 |
| 45 | H11 | -1.187 | -0.858 | 2.091 | HA3 | 1 | HEM | 0.09 |
| 46 | H12 | -1.042 | -0.926 | 2.123 | HA3 | 1 | HEM | 0.09 |
| 47 | C28 | -0.991 | -0.794 | 1.764 | CE1 | 1 | HEM | -0.15 |
| 48 | H13 | -0.984 | -0.778 | 1.665 | HE1 | 1 | HEM | 0.15 |
| 49 | C29 | -0.937 | -0.706 | 1.842 | CE2 | 1 | HEM | -0.42 |
| 50 | H14 | -0.941 | -0.719 | 1.941 | HE2 | 1 | HEM | 0.21 |
| 51 | H15 | -0.890 | -0.627 | 1.804 | HE2 | 1 | HEM | 0.21 |
| 52 | C30 | -1.013 | -1.001 | 1.289 | CT3 | 1 | HEM | -0.27 |
| 53 | H16 | -1.014 | -1.023 | 1.191 | HA3 | 1 | HEM | 0.09 |
| 54 | H17 | -1.058 | -0.913 | 1.304 | HA3 | 1 | HEM | 0.09 |
| 55 | H18 | -0.919 | -0.996 | 1.321 | HA3 | 1 | HEM | 0.09 |
| 56 | C31 | -1.153 | -1.267 | 1.178 | CE1 | 1 | HEM | -0.15 |
| 57 | H19 | -1.217 | -1.343 | 1.161 | HE1 | 1 | HEM | 0.15 |
| 58 | C32 | -1.084 | -1.224 | 1.078 | CE2 | 1 | HEM | -0.42 |
| 59 | H20 | -1.019 | -1.148 | 1.091 | HE2 | 1 | HEM | 0.21 |
| 60 | H21 | -1.095 | -1.265 | 0.987 | HE2 | 1 | HEM | 0.21 |
| 61 | C33 | -1.399 | -1.669 | 1.371 | CT3 | 1 | HEM | -0.27 |
| 62 | H22 | -1.442 | -1.757 | 1.391 | HA3 | 1 | HEM | 0.09 |
| 63 | H23 | -1.462 | -1.611 | 1.320 | HA3 | 1 | HEM | 0.09 |

| | | | | | | | | |
|---|---|---|---|---|---|---|---|---|
| 64 | H24 | -1.316 | -1.684 | 1.317 | HA3 | 1 | HEM | 0.09 |
| 65 | C34 | -1.451 | -1.778 | 1.650 | CT2 | 1 | HEM | -0.18 |
| 66 | H25 | -1.524 | -1.836 | 1.615 | HA2 | 1 | HEM | 0.09 |
| 67 | H26 | -1.470 | -1.752 | 1.744 | HA2 | 1 | HEM | 0.09 |
| 68 | C35 | -1.356 | -1.889 | 1.682 | CT2 | 1 | HEM | -0.28 |
| 69 | H27 | -1.292 | -1.858 | 1.753 | HA2 | 1 | HEM | 0.09 |
| 70 | H28 | -1.305 | -1.913 | 1.599 | HA2 | 1 | HEM | 0.09 |
| 71 | C36 | -1.424 | -2.012 | 1.731 | CC | 1 | HEM | 0.62 |
| 72 | O3 | -1.533 | -2.013 | 1.783 | OC | 1 | HEM | -0.76 |
| 73 | O4 | -1.369 | -2.116 | 1.721 | OC | 1 | HEM | -0.76 |

**Table S5.** Parameters of bond, angle and torsion for cofactors.

| Cofactor | improper type | *Vn*/2 (kcal/mole) | γ(degrees) | n | |
|---|---|---|---|---|---|
| riboflavin | CG2O6-NG2D1-NG2S1-OG2D1 | 669.440000 | 0.000000 | 2 | General improper torsional angle (2 general atom types) |
| NADPH | CG2D1O-CG2D1-NG3P1-HGA4 | 443.504000 | 0.000000 | 2 | General improper torsional angle (2 general atom types) |
| | CG2D1O-CG2DC1-NG3P1-HGA4 | 443.504000 | 0.000000 | 2 | |

**Table S6.** The primers for site-directed mutagenesis.

| variant | primer |
|---|---|
| S56A-F | 5’-CGCACTGAACACCTTCGGGTTGCTCAGGAT-3’ |
| S56A-R | 5’-GATACCATGCGTTTAGATCCGATCAAACT-3’ |
| D74A-F | 5’- CGCGCCTTCCACAATGGCCTCGTCCAGT-3’ |
| D74A-R | 5’- TTTGCCCACACAGACCCTCCGAAACAC-3’ |
| M159A-F | 5’- CGCCTTGTCCATCCACTGGCGAAACAG-3’ |
| M159A-R | 5’- CTGGACGGTAGCGAAAAGTTCGAAAGC-3’ |
| L160A-F | 5’- CGCCATCTTGTCCATCCACTGGCGAAACAG-3’ |
| L160A-R | 5’- GACGGTAGCGAAAAGTTCGAAAGCCCGGAAAC-3’ |
| G162A-F | 5’- CGCGTCCAGCATCTTGTCCATCCACTGG-3’ |
| G162A-R | 5’- AGCGAAAAGTTCGAAAGCCCGGAAACCGT-3’ |
| F166A-F | 5’- CGCCTTTTCGCTACCGTCCAGCAT-3’ |
| F166A-R | 5’- GAAAGCCCGGAAACCGTGCTGGAAC-3’ |
| R233A-F | 5’- CGCGTTGGCGATATTACTAATCTGG-3’ |
| R233A-R | 5’- TTACTGGTGAACGGTCATCTGACAAC-3’ |
| V236A-F | 5’- CGCCAGTAAGCGGTTGGCGATATTACTAAT-3’ |
| V236A-R | 5’- AACGGTCATCTGACAACCGCCATGCTGAT-3’ |
| N237A-F | 5’- CGCCACCAGTAAGCGGTTGGCGAT-3’ |
| N237A-R | 5’- GGTCATCTGACAACCGCCATGCTGAT-3’ |
| T241A-F | 5’- CGCCAGATGACCGTTCACCAGTAAGCGGT-3’ |
| T241A-R | 5’- ACCGCCATGCTGATCGCCAATACCAT-3’ |
| R289A-F | 5’- CGCGCCCACGCCACAAATCGGGCTCATGT-3’ |
| R289A-R | 5’- GCAACAAATAGCGAAGTGGAAGTGG-3’ |
| L340F-F | 5’-ACCCAGATGGGCATTCGGGCTACGGCCTGCGTCAAA AACAT-3’ |
| L340F- R | 5’-TTTGGCCGTGGCATTCATTTTTGCCTGGGCCGTCAGC T-3’ |
| L340F&C346A-F | 5’-CGCAAAATGAATGCCACGGCCAAAAC-3’ |
| C347A-F | 5’- CGCAAAATGAATGCCACGGCCTAAAC-3’ |
| C347A-R | 5’- CTGGGCCGTCAGCTGGCACGCATGG-‘ |
| D386A-F | 5’- CGCCACCACCTGCAGAAAGGTCGGCGGAT-3’ |
| D386A-R | 5’- GCCAGCGGTGTTGCCACCCTGCCTGT-3’ |

| | |
|---|---|
| R190A-F | 5’- CGCCATCTCCCACAGCAGTTCCAGTTCTTTAT-3’ |
| R190A-R | 5’- GACTATTGGCACGAACGCGCAGCAGAGAGT-3’ |
| G288A-F | 5’- CGCCACGCCACAAATCGGGCTCATGTAACGCAT-3’ |
| G288A-R | 5’- CGCGCAACAAATAGCGAAGTGGAAGTGGCCGGT-3’ |

Bold CGC is an optimized codon of alanine. Bold TTT or AAA is an optimized codon of Phenylalanine.

### 3. Supplementary Figures

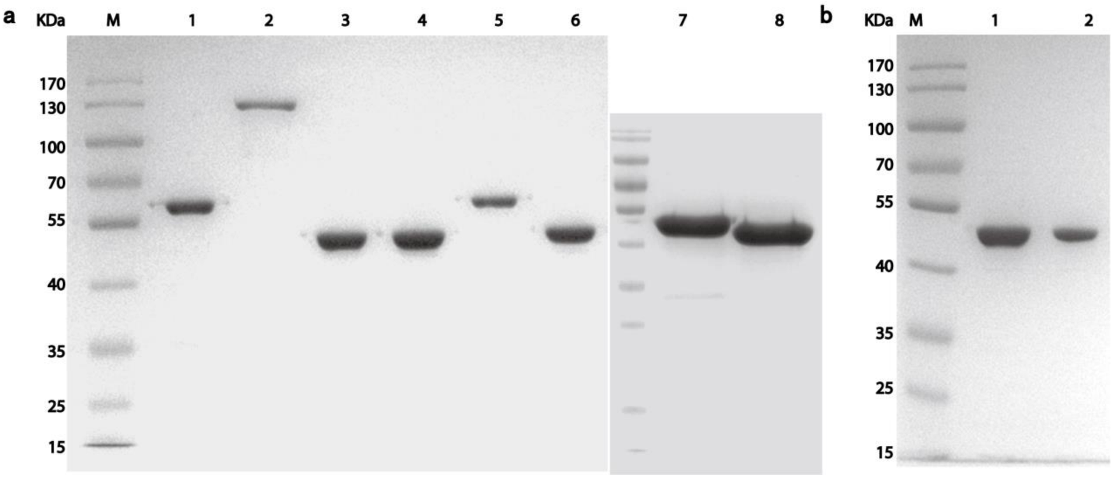

**Figure S1.** PAGE analyses of purified recombinant microbial P450 proteins.

(a) SDS-PAGE was performed on a 10% gel under reduced condition. M, molecular weight marker; Lane 1, BMP; Lane 2, P450 BM3; Lane 3, HmtS; Lane 4, HmtS-L340F; Lane 5, HmtN; Lane 6, HmtT; Lane 7, CYP106A2; Lane 8, CYP107DY1;

(b) Non-reduced PAGE was performed on a 10% gel on iced water. M, molecular weight marker; Lane 1, HmtS-L340F; Lane 2, HmtS-L340F in the presence of NADPH, riboflavin and diphenhydramine.

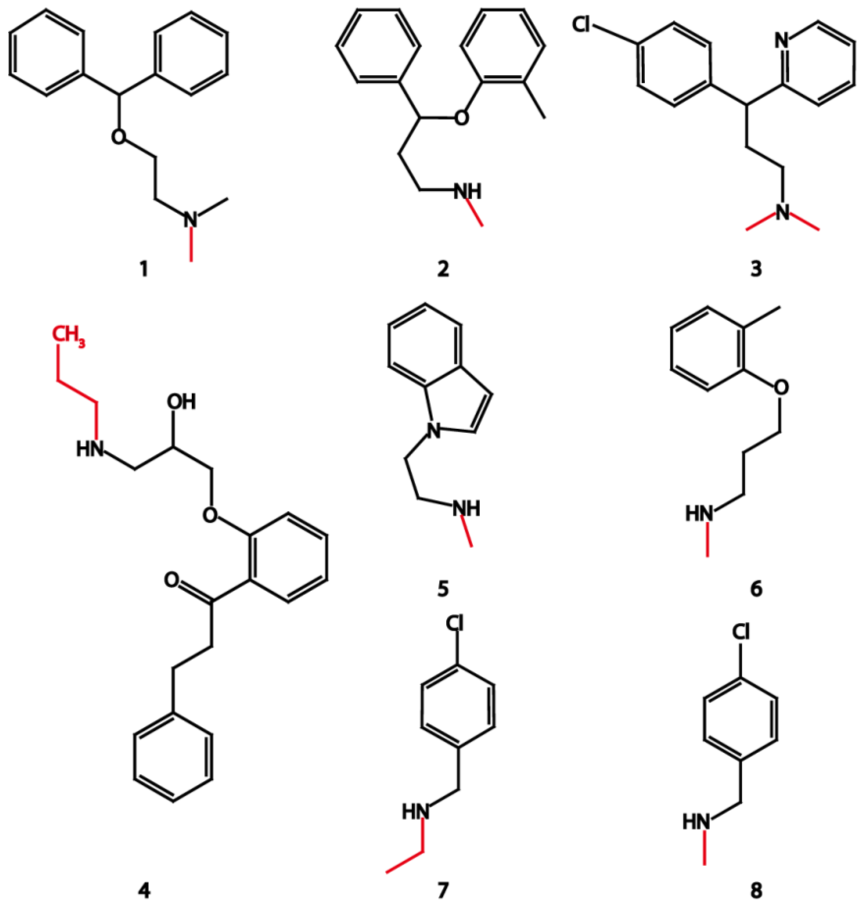

**Figure S2.** Chemical structures of the substrates for the screening of HmtS with riboflavin.

The group with red color in the structures indicates the alkyl groups to be oxidatively cleaved.

**1**: 2-(benzhydryloxy)-N,N-dimethylethanamine (diphenhydramine);

**2**: N-methyl-3-phenyl-3-(o-tolyloxy)propan-1-amine;

**3**: 3-(4-chlorophenyl)-N,N-dimethyl-3-(pyridin-2-yl)propan-1-amine;

**4**: 1-(2-(2-hydroxy-3-(propylamino)propoxy)phenyl)-3-phenylpropan-1-one;

**5**: 2-(1H-indol-1-yl)-N-methylethanamine;

**6**: N-methyl-3-(o-tolyloxy)propan-1-amine;

**7**: N-(4-chlorobenzyl) ethanamine;

**8**: 1-(4-chlorophenyl)-N-methylmethanamine.

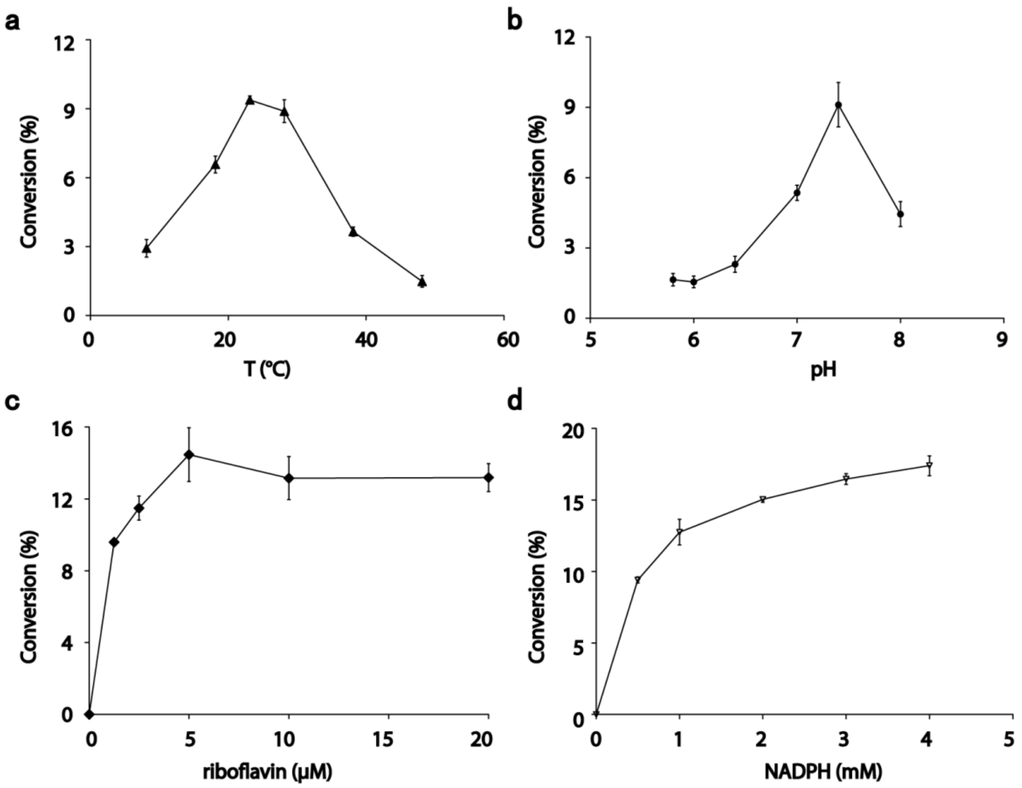

**Figure S3.** Catalytic characteristics of HmtS-riboflavin system.

(a) Temperature optima of the catalytic system. (b) pH optima of the catalytic system. (c) The effects of riboflavin concentration on the catalytic system. (d) The effects of NADPH concentration on the catalytic system. Data are average values of 3 independent measurements with standard deviations. The reactions for HmtS-riboflavin system were performed for 4 h using 4 μM enzyme, 0-4 mM NADPH, 340 μM diphenhydramine and 0-20 μM riboflavin in buffer. Temperatures ranging from 15 ℃ to 55 ℃ were used for the reactions. Optimal pH was examined from pH 5.8-8.0.

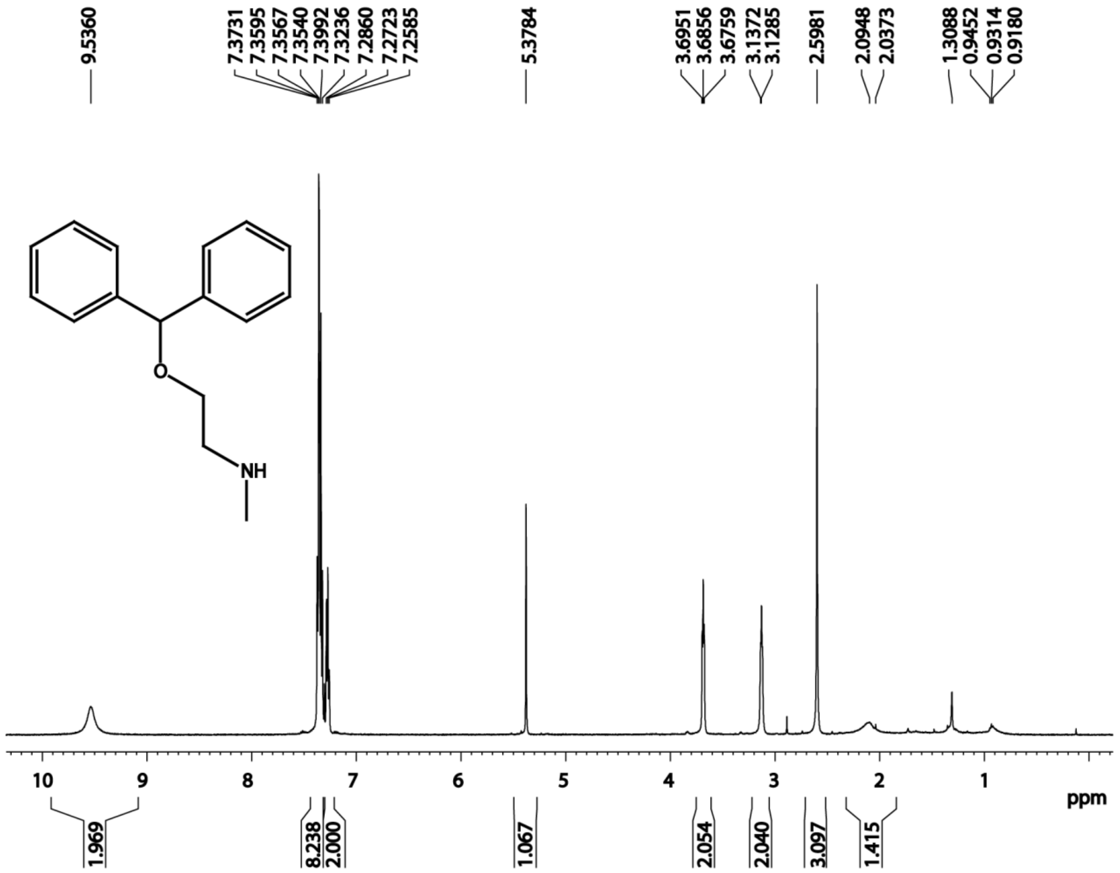

**Figure S4.** $^{1}$H NMR of the N-dealkylation product 2-(benzhydryloxy)-N-methylethanamine.

$^{1}$H NMR (500 MHz, $CDCl_3$) δ 9.53 (s, 1H) 7.32-7.37 (m, 8H),7.27-7.30 (m,2H),5.38 (s, 1H), 3.69 (t, $J$=5 Hz, 2H), 3.13 (t, $J$ = 5.00 Hz, 2H), 2.60 (s, 3H).

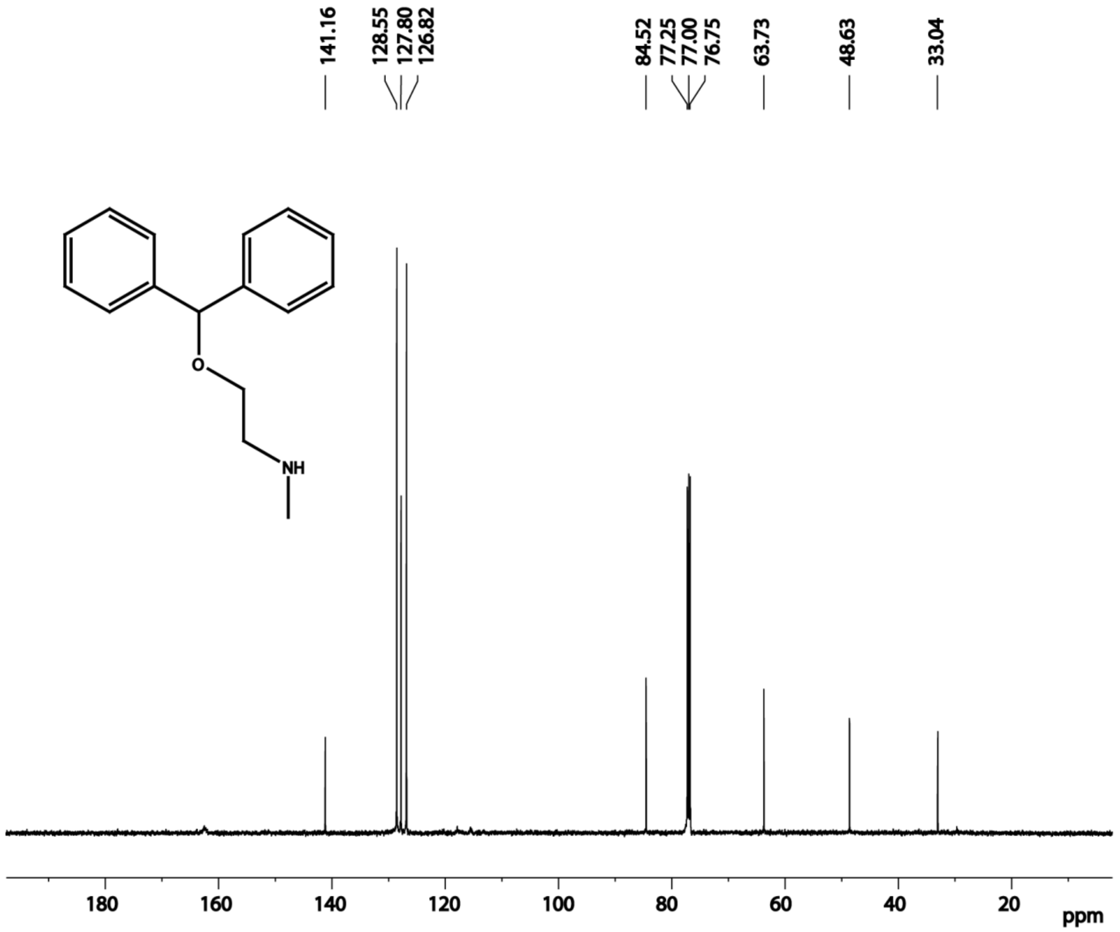

**Figure S5.** $^{13}$C NMR of the N-dealkylation product 2-(benzhydryloxy)-N-methylethanamine.

$^{13}$C NMR (125 MHz, $CDCl_3$) δ 141.16 (C), 141.16 (C), 128.55 (CH), 128.55 (CH), 128.55 (CH), 128.55 (CH), 127.8 (CH), 127.8 (CH), 127.8 (CH), 127.8 (CH), 126.82 (CH), 126.82 (CH), 84.5 (CH), 63.73 ($CH_2$), 48.63 ($CH_2$), 33.04 ($CH_3$).

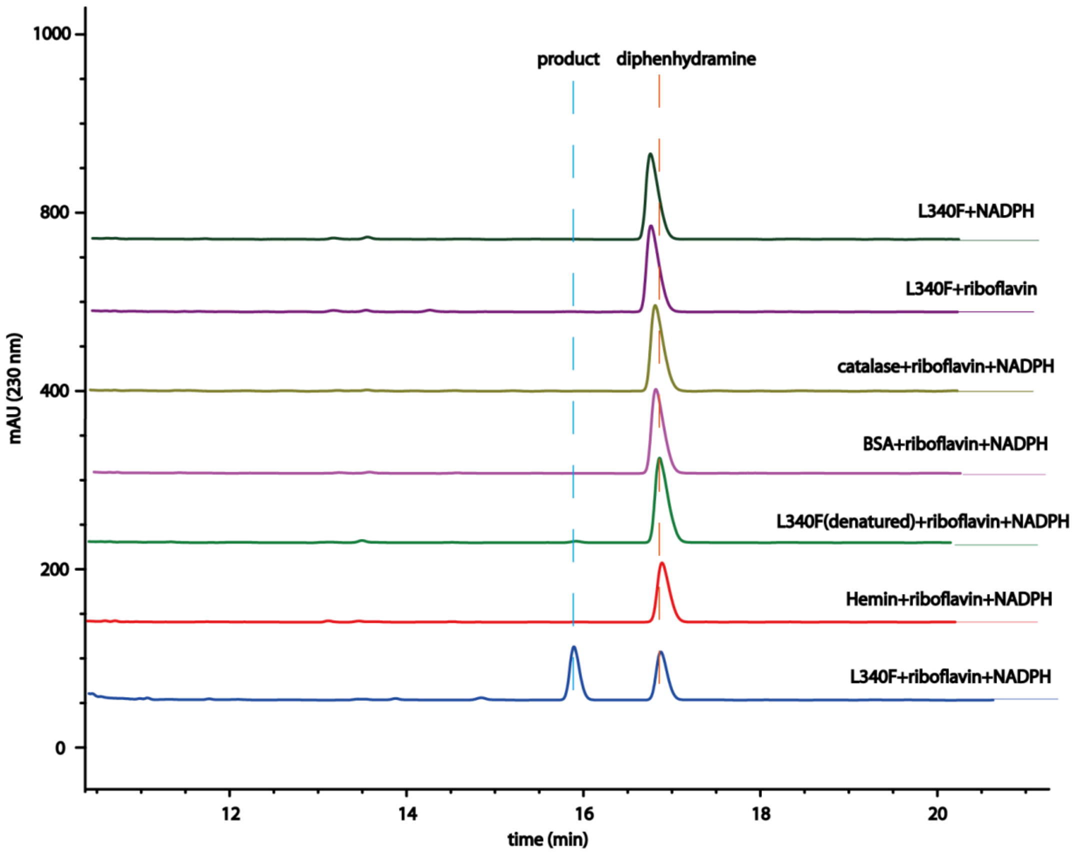

**Figure S6.** HPLC chromatograms of riboflavin mediated N-dealkylation of diphenhydramine by HmtS-L340F. The reaction mixtures contained 4 μM purified HmtS-L340F, catalase, hemin or BSA, 400 μM substrate, 3 mM NADPH, 4 μM riboflavin in PBS buffer (pH7.4) and were conducted for 4 h.

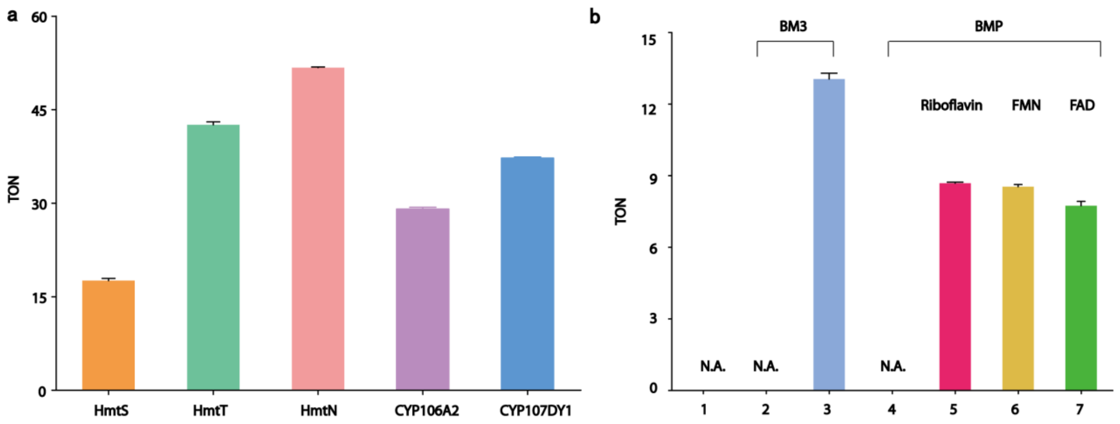

**Figure S7.** Comparison of different microbial P450 enzymes on the N-dealkylation of diphenhydramine.

(a) Comparison of five different microbial P450 enzymes on the N-dealkylation of diphenhydramine. (b) Comparison of the catalytic efficiency for BM3 and BMP. 1: NADPH; 2: BM3; 3: NADPH + BM3; 4: BMP + NADPH; 5: BMP + riboflavin + NADPH; 6: BMP + FMN + NADPH; 7: BMP + FAD + NADPH.

Data are average values of 3 independent measurements with standard deviations. The reaction mixtures contained 4 μM purified enzyme, 400 μM substrate, 3 mM NADPH, 4 μM riboflavin in PBS buffer (pH7.4) and were conducted for 4 h.

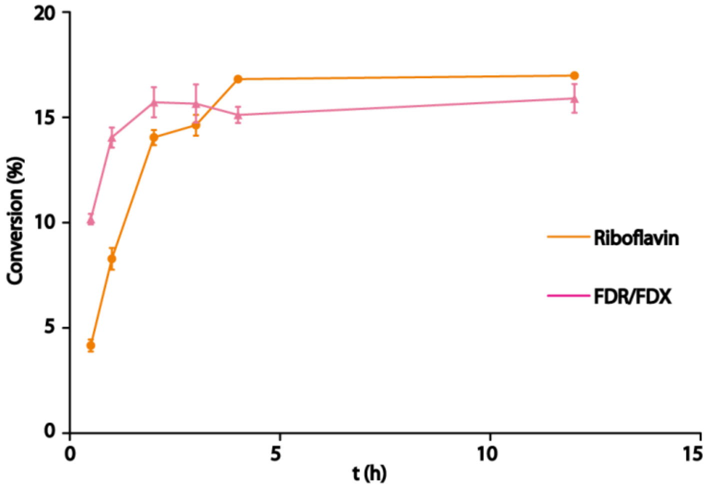

**Figure S8.** The initial rate of N-dealkylation of diphenhydramine catalyzed by HmtS in the presence of riboflavin or FDR/FDX. Data are average values of 3 independent measurements with standard deviations. The reactions for FDR/FDX system contained 4 μM purified enzyme, 340 μM substrate, 3 mM NADPH, 5 μM spinach ferredoxin $NADP^+$ reductase, 1 μM spinach ferredoxin in PBS buffer (pH7.4) and were conducted for 0.5-12 h. The reactions for riboflavin system contained 4 μM purified enzyme, 340 μM substrate, 3 mM NADPH, 4 μM riboflavin in PBS buffer (pH7.4) and were conducted for 0.5-12 h.

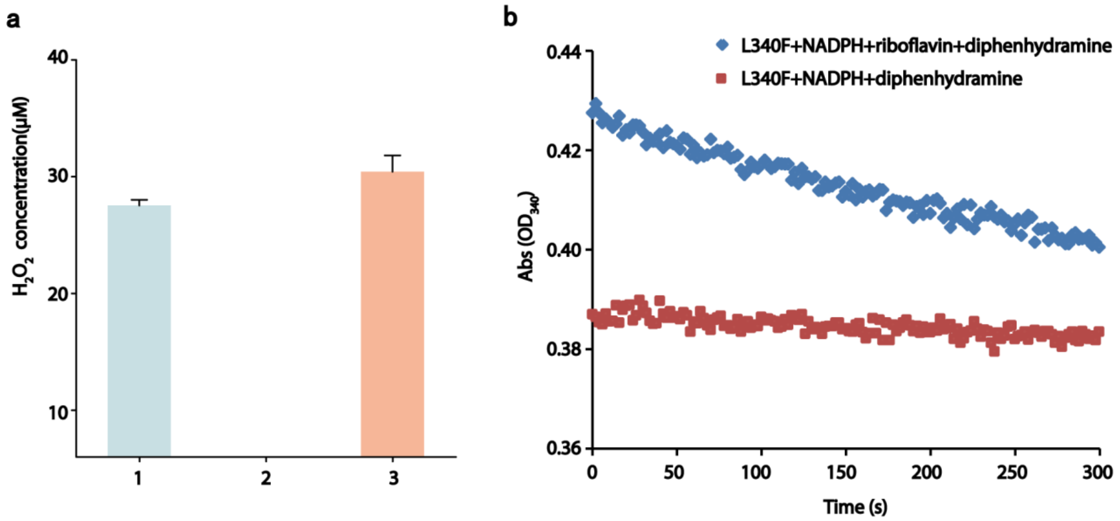

**Figure S9.** Comparison of different reactions on $H_2O_2$ generation.

Data are average values of 3 independent measurements with standard deviations.

(a) The assay was carried out in PBS buffer pH 7.4 (final volume 0.8 ml) with 0.5 ml reaction mixture containing 125 μM phenol, 1.25mM 4-aminoantipyrine and 0.1 mg/ml horseradish peroxidase for 45 min. 1: FDR/FDX + NADPH + diphenhydramine; 2: riboflavin + NADPH + diphenhydramine; 3: HmtT + NADPH + riboflavin. (b) The reaction mixtures for NADPH consumption contained 2 μM purified enzyme, 100 μM substrate, 50 μM NADPH, 3 μM riboflavin in PBS buffer (pH7.4), and the absorbance at 340 nm was measured for 300 s.

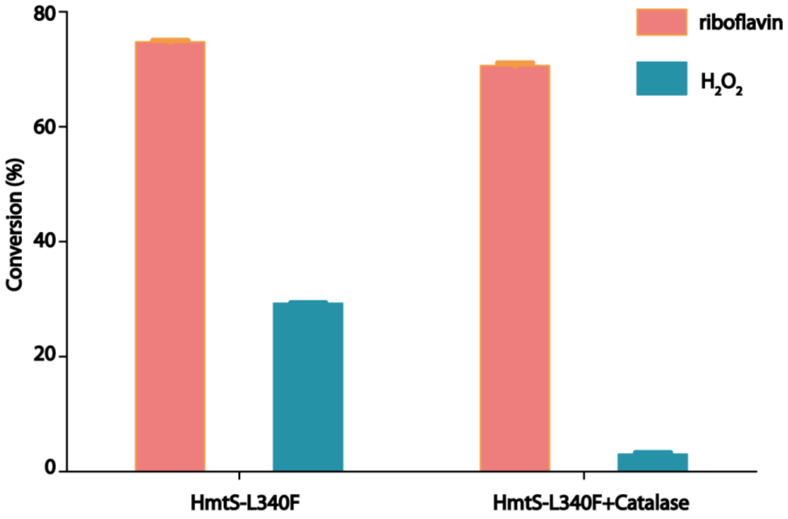

**Figure S10.** Comparison of riboflavin and $H_2O_2$ on the dealkylation of diphenhydramine by HmtS-L340F.

The reactions with riboflavin contained 4 µM purified HmtS-L340F , 400 µM substrate, 3 mM NADPH, 4 µM riboflavin in PBS buffer (pH7.4) and were conducted for 4 h. The reactions with riboflavin and catalase contained 1U/ml catalase, 400 µM substrate, 3 mM NADPH, 4 µM riboflavin, 4 µM HmtS-L340F in PBS buffer (pH7.4) and were conducted for 4 h. The reactions with $H_2O_2$ contained 4 µM purified HmtS-L340F, 400 µM substrate, 1 mM $H_2O_2$ in PBS buffer (pH7.4) and were conducted for 4 h. The reactions for $H_2O_2$ and catalase contained 1 U/ml catalase, 4 µM HmtS-L340F, 400 µM substrate, 1 mM $H_2O_2$ in PBS buffer (pH7.4) and were conducted for 4 h.

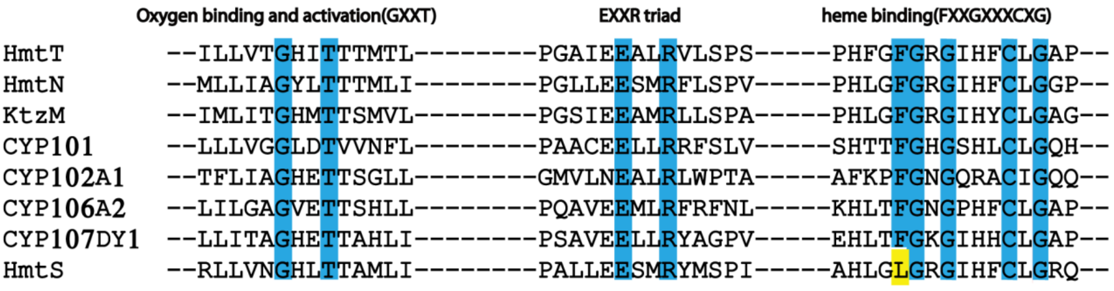

**Figure S11.** Sequence alignments of HmtS with P450-family enzymes listed in Table S2.

ClustalX was used for the alignments. Conserved residues are indicated in blue. The different residue of HmtS in the heme binding motif is highlighted in yellow.

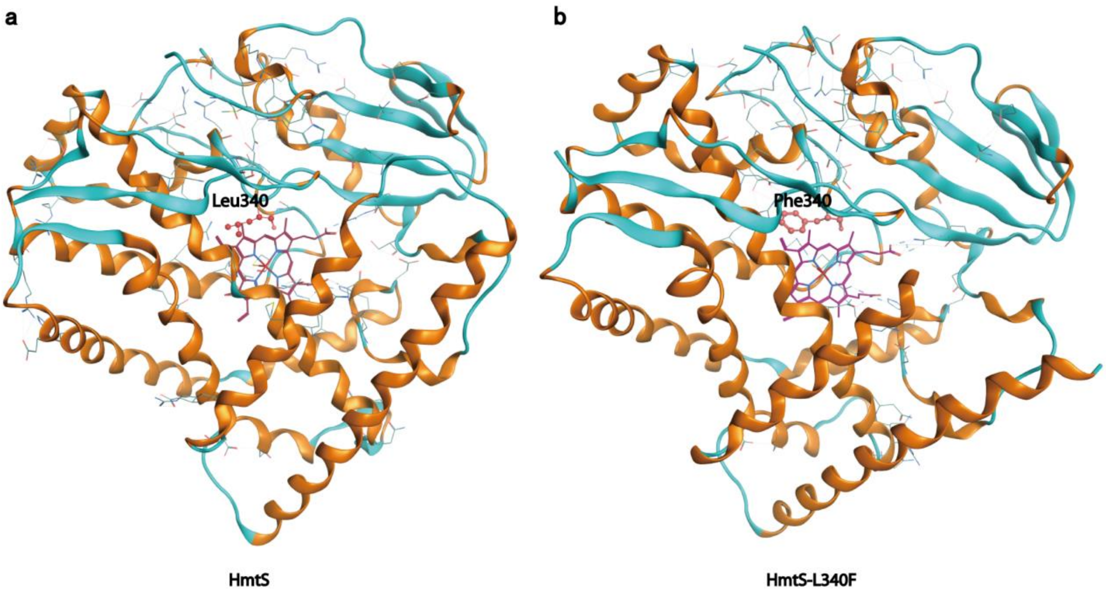

**Figure S12.** Crystal structures of HmtS and HmtS-L340F mutant.

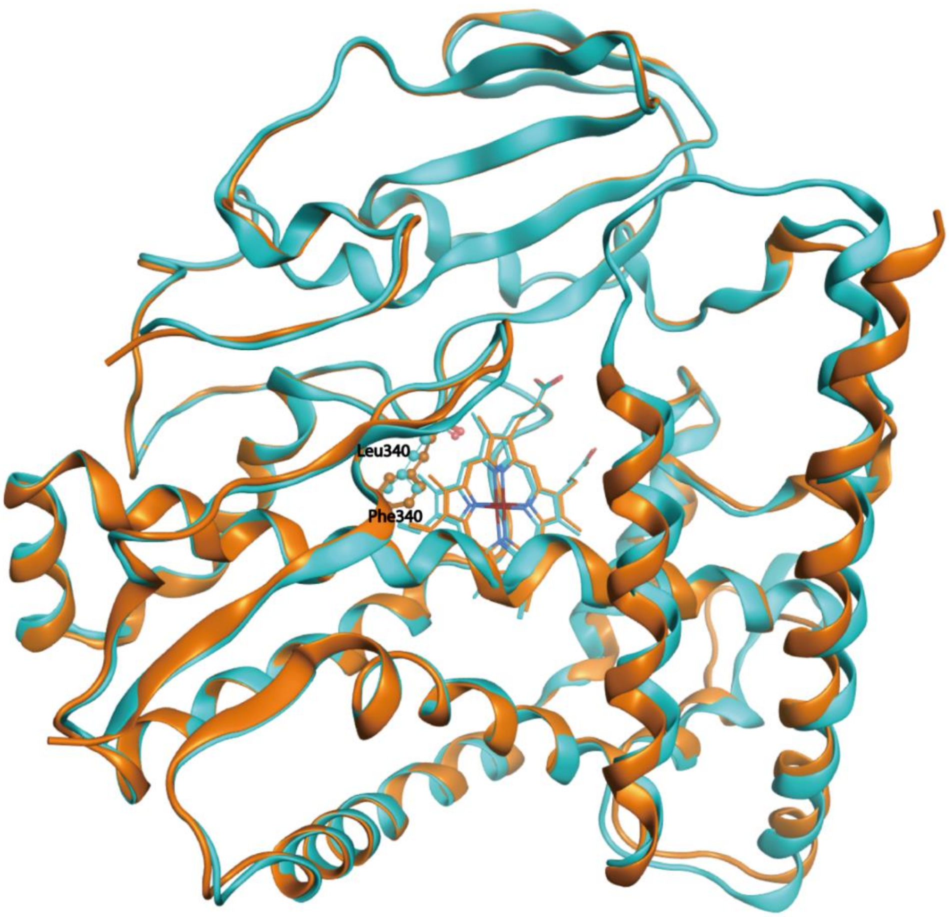

**Figure S13.** Superimposition of the structures of HmtS (PDB: 5Z9I) and HmtS-L340F mutant (PDB: 5Z9J). HmtS is in blue; HmtS-L340F mutant is in orange.

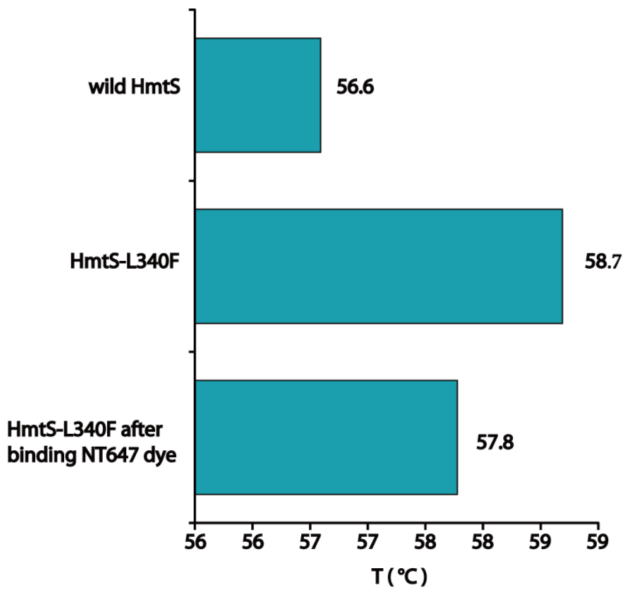

**Figure S14**. Thermal stability of HmtS and HmtS-L340F mutant. Data are average values of 3 independent measurements.

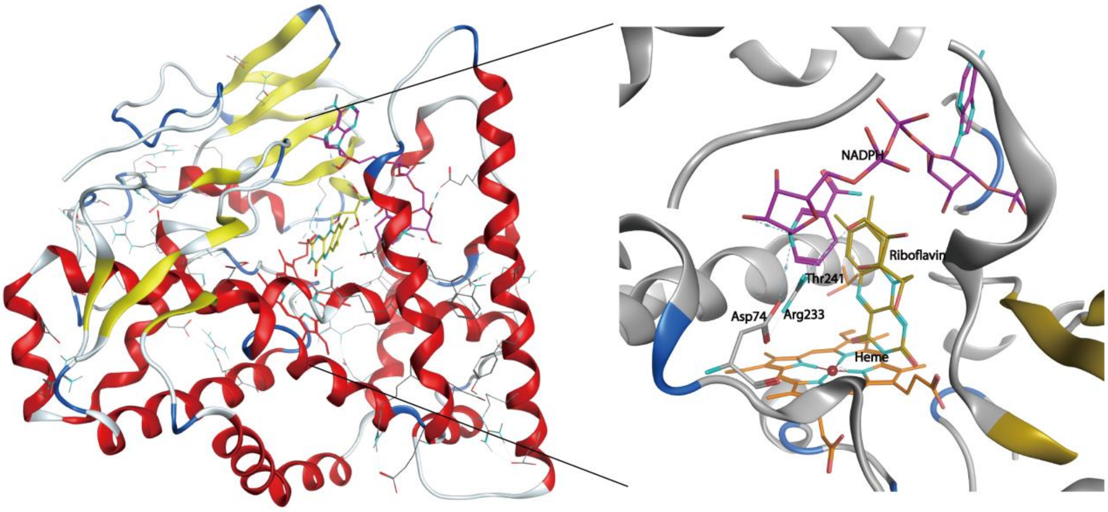

**Figure S15.** Modeled complex structure of NADPH and riboflavin with HmtS-L340F. Ribbon diagram of NADPH (violet) and riboflavin (yellow) is shown in the model.

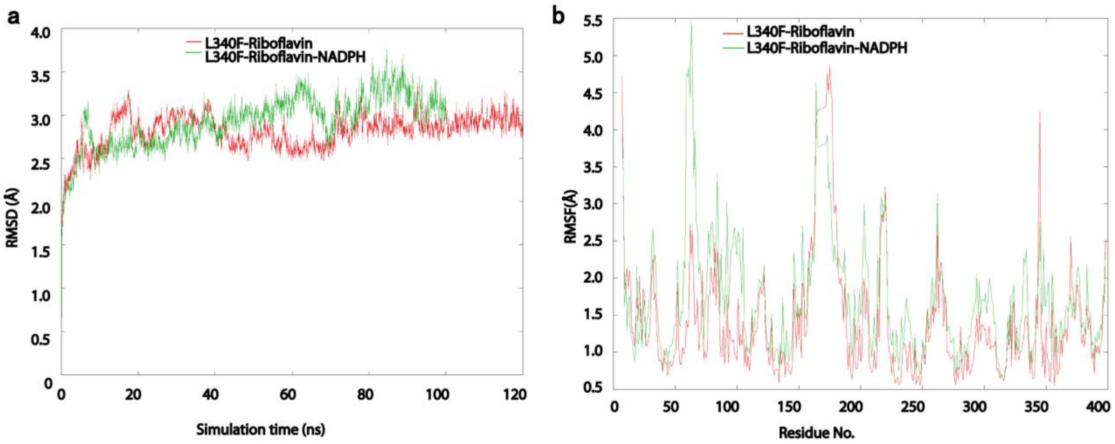

**Figure S16.** MD simulation of the modeled structures of HmtS-L340F complexed with riboflavin and HmtS-L340F complexed with riboflavin and NADPH.

(a) RMSD of HmtS-L340F as a function of time; (b) RMSF of all Cα atoms from the crystal structure as the template.

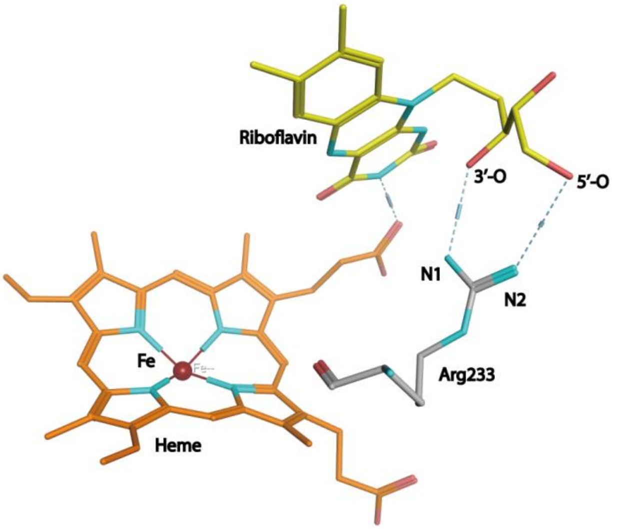

**Figure S17.** Snapshot of the simulation of HmtS-L340F-riboflavin complex for the interactions between riboflavin and residue Arg233.

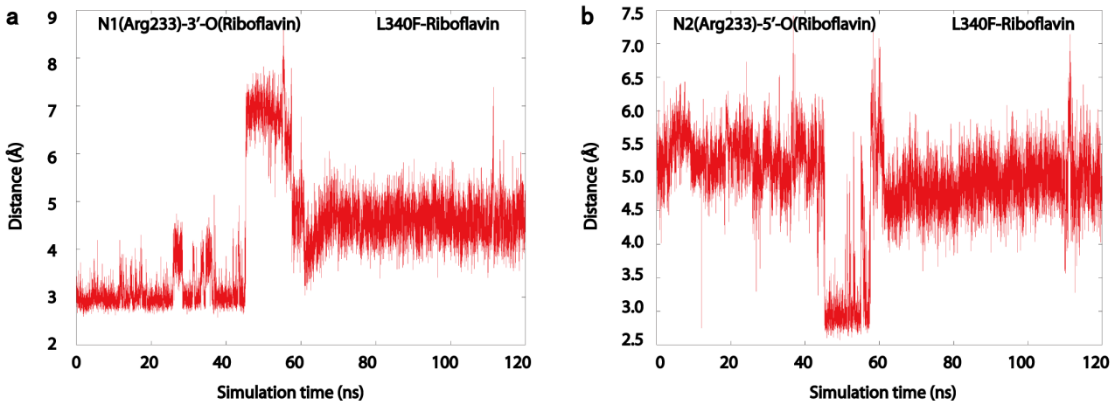

**Figure S18.** MD simulation of HmtS-L340F-riboflavin complex. Distance fluctuations of Arg233 to riboflavin.

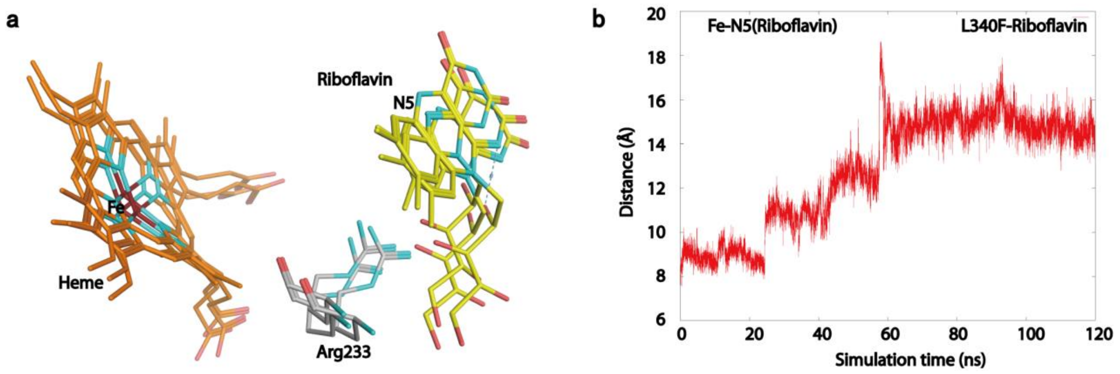

**Figure S19.** Serial spatial positions between riboflavin and heme in HmtS-L340F-riboflavin complex. Ribbon diagram of riboflavin (yellow) is shown in the models.

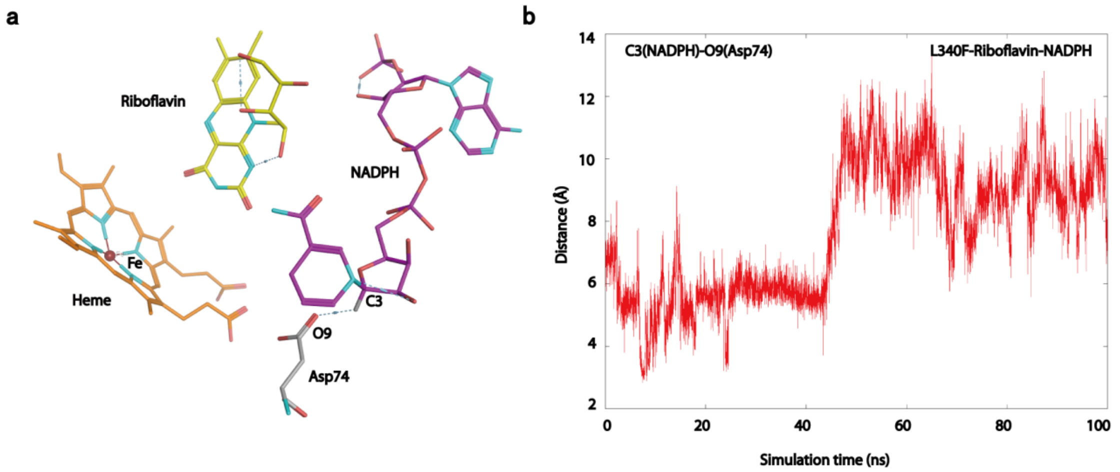

**Figure S20.** Binding mode of HmtS-L340F with NADPH and riboflavin.

(a) Snapshot of the stimulation of HmtS-L340F-riboflavin-NADPH for the interactions between NADPH with residue Asp74; (b) Distance fluctuations of Asp74 to NADPH in HmtS-L340F-riboflavin-NADPH complex.

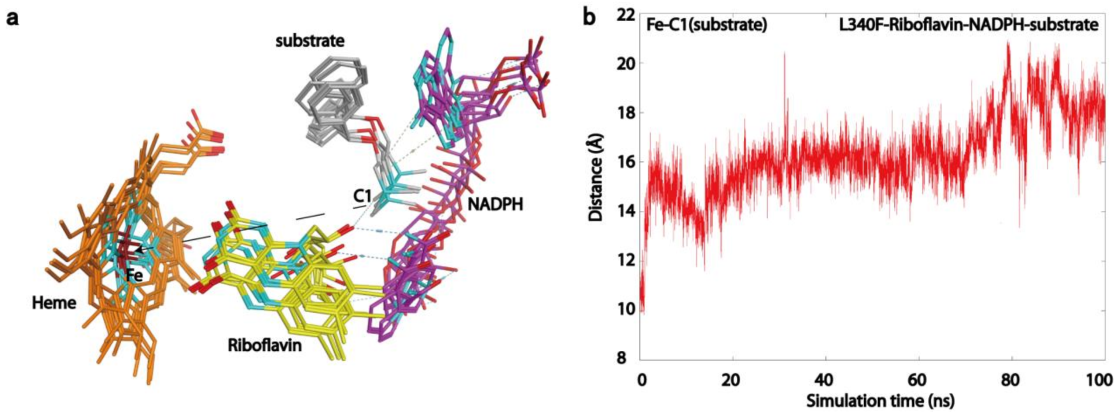

**Figure S21**. Serial spatial positions between substrate and heme in HmtS-L340F-riboflavin-NADPH-substrate complex. Ribbon diagram of riboflavin (yellow), NADPH (violet) and substrate (gray) is shown in the models.

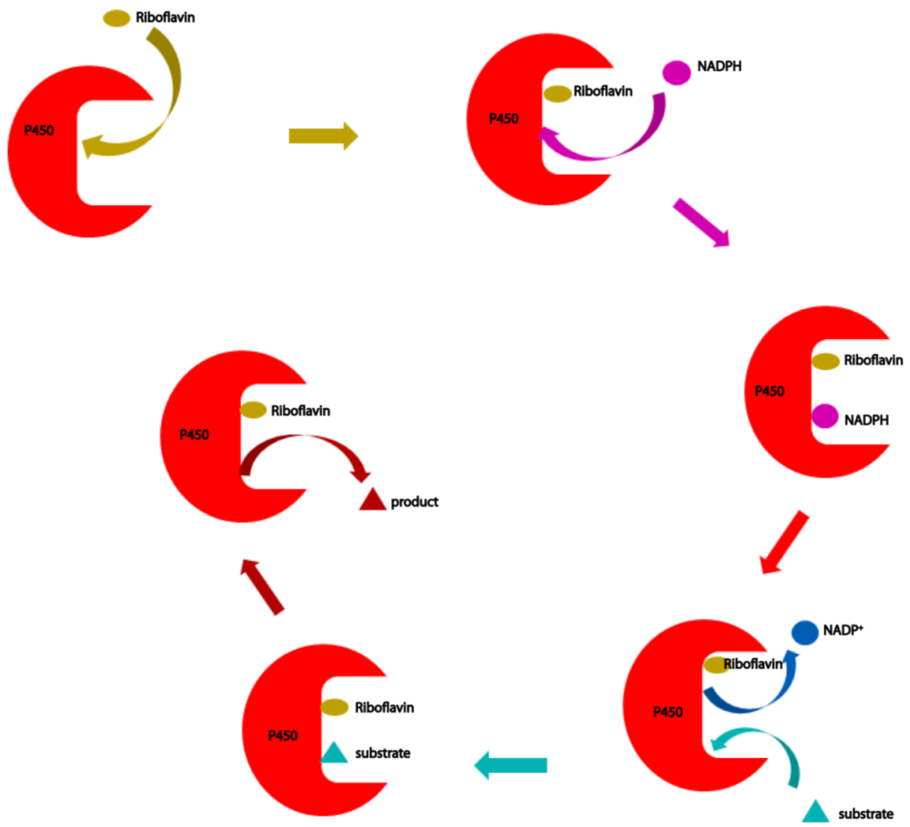

**Figure S22.** Proposed sequential model for the reaction mediated by riboflavin and NADPH for microbial P450 monooxygenases.

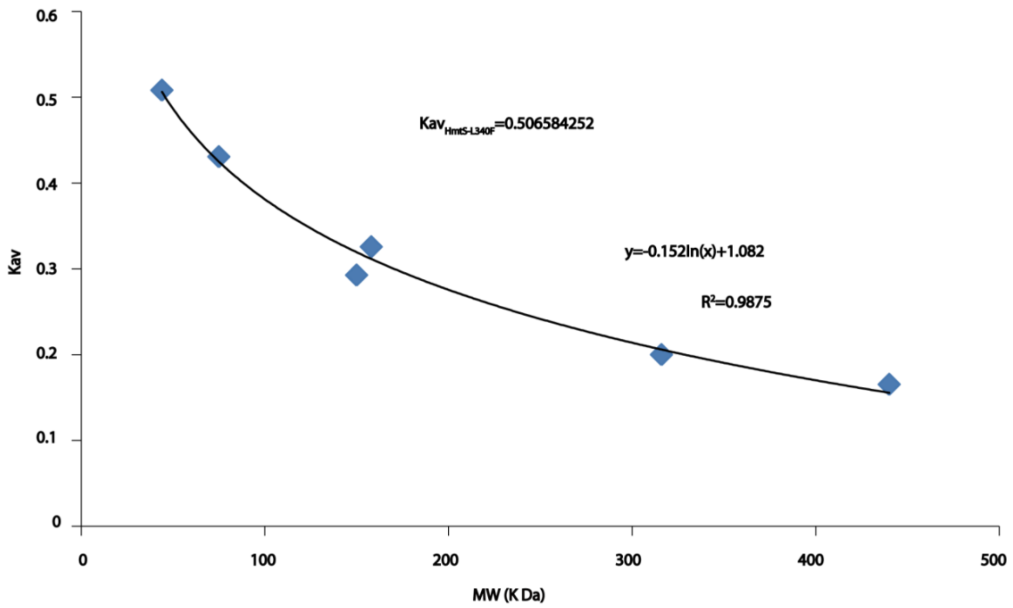

**Figure S23.** Molecular weight determination of HmtS-L340F by gel filtration chromatography. Sephacryl S-200HR was used and calibrated with standard proteins. The molecular weight of HmtS-L340F was determined to be 44.06 kDa.

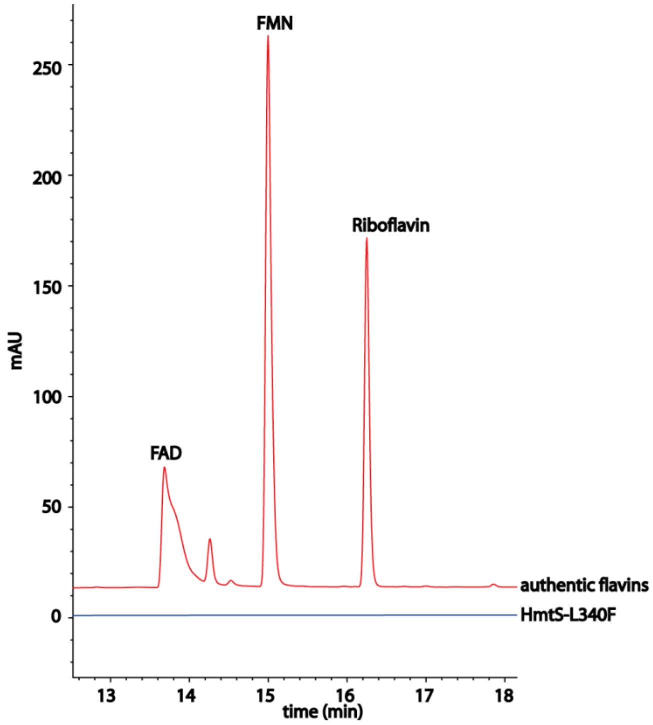

**Figure S24**. HPLC determination of bound flavins in HmtS-L340F. Authentic flavins are in red; HPLC trace for 3 mg/ml HmtS-L340F after boiling is in blue.

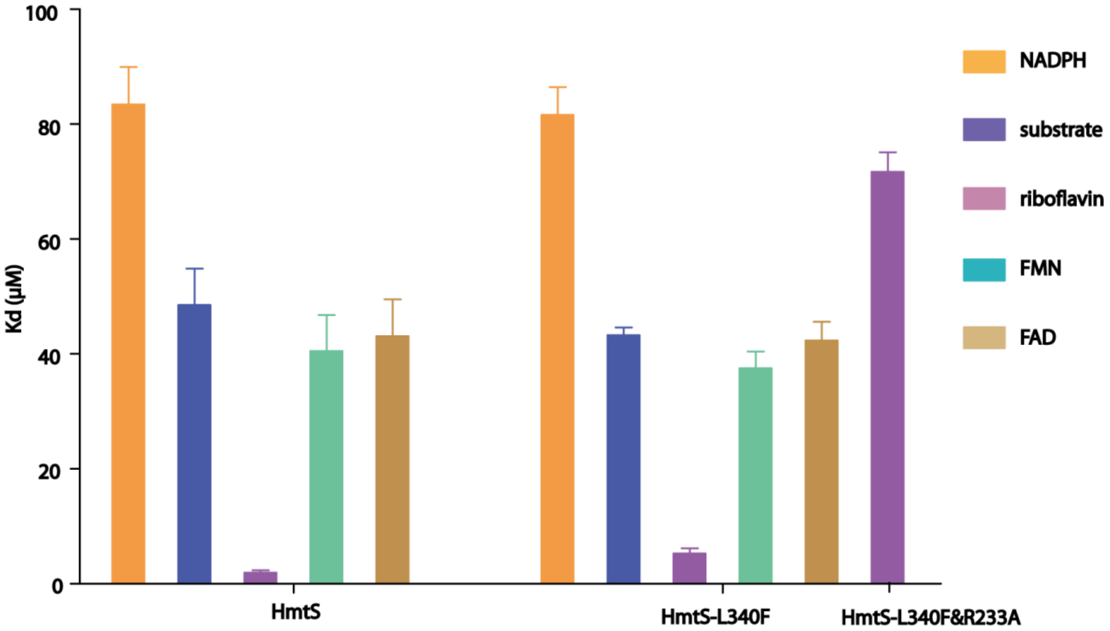

**Figure S25.** Binding affinity determined by Microscale Thermophoresis (MST).

$K_d$ values were calculated by MO Affinity Analysis v2.2.4. $K_d$ valu~~s~~es for the binding of NADPH, diphenhydramine, riboflavin, FMN and FAD with HmtS were determined and calculated to be 83 µM, 35 µM, 1.9 µM, 40 µM and 43 µM, respectively. $K_d$ values for the binding of NADPH, diphenhydramine, riboflavin, FMN and FAD with HmtS-L340F were determined and calculated to be 82 µM, 43 µM, 5.3 µM, 38 µM and 42 µM, respectively. *Kd* values for the binding of riboflavin with HmtS-L340F &R233A were 71 µM.

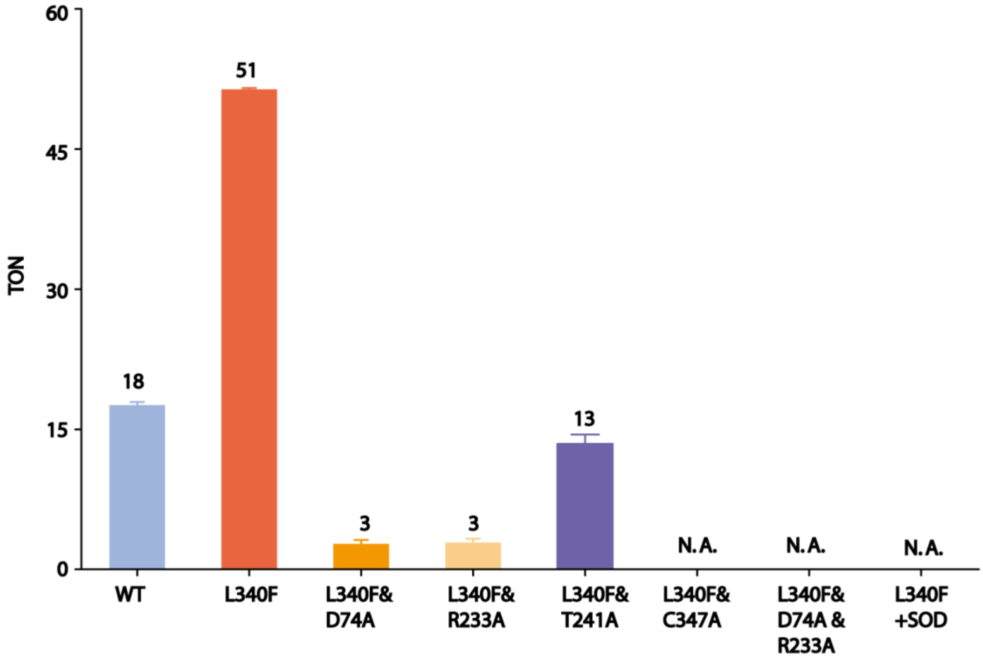

**Figure S26.** Comparison of different mutants of HmtS and additional SOD on the catalytic efficiency.

Data are average values of 3 independent determinations. All reactions were conducted under the same conditions. N.A. represents not active. The reaction mixtures contained 4 μM purified enzyme, 400 μM substrate, 3 mM NADPH, 4 μM riboflavin in PBS buffer (pH7.4) and were conducted for 4 h. Additional SOD was added into the reaction mixtures at a final concentration of 1 U/ml.
